# Enhanced detection of RNA modifications and mappability with high-accuracy nanopore RNA basecalling models

**DOI:** 10.1101/2023.11.28.568965

**Authors:** Gregor Diensthuber, Leszek Pryszcz, Laia Llovera, Morghan C Lucas, Anna Delgado-Tejedor, Sonia Cruciani, Jean-Yves Roignant, Oguzhan Begik, Eva Maria Novoa

**Author notes:** These authors contributed equally (Gregor Diensthuber, Leszek Pryszcz). Correspondence to: Oguzhan Begik and Eva Maria Novoa.

## Abstract

In recent years, nanopore direct RNA sequencing (DRS) has established itself as a valuable tool for studying the epitranscriptome, due to its ability to detect multiple modifications within the same full-length native RNA molecules. While RNA modifications can be identified in the form of systematic basecalling ‘errors’ in DRS datasets, *N6*-methyladenosine (m^6^A) modifications produce relatively low ‘errors’ compared to other RNA modifications, limiting the applicability of this approach to m^6^A sites that are modified at high stoichiometries. Here, we demonstrate that the use of alternative RNA basecalling models, trained with fully unmodified sequences, increases the ‘error’ signal of m^6^A, leading to enhanced detection and improved sensitivity even at low stoichiometries. Moreover, we find that high-accuracy alternative RNA basecalling models can show up to 97% median basecalling accuracy, outperforming currently available RNA basecalling models, which show 91% median basecalling accuracy. Notably, the use of high-accuracy basecalling models is accompanied by a significant increase in the number of mapped reads –especially in shorter RNA fractions– and increased basecalling error signatures at pseudouridine (Ψ) and N1-methylpseudouridine (m^1^Ψ) modified sites. Overall, our work demonstrates that alternative RNA basecalling models can be used to improve the detection of RNA modifications, read mappability and basecalling accuracy in nanopore DRS datasets.

## INTRODUCTION

RNA modifications play a crucial role in a wide variety of biological processes, including gene expression regulation, RNA stability, splicing and translation (1–3). Our understanding of RNA modifications has vastly increased in the past years with the development of novel methods that can chart the modification landscape in a transcriptome-wide fashion (4–6). Next-generation sequencing (NGS)-based methods were the first to revolutionize the field of RNA modification detection by providing high-accuracy maps for *N6*-methyladenosine (m^6^A), 2′-O-methylations (Nm), *N4*-acetylcytosine (ac^4^C) and pseudouridine (Ψ), among others (7–13). More recently, nanopore direct RNA sequencing (DRS) has emerged as a promising alternative to NGS for RNA modification detection due to its ability to directly detect RNA modifications in native RNA reads without the need of additional chemical labeling steps (14–17) and in full-length RNA reads, thus permitting isoform-specific RNA modification analyses.

Nanopore direct RNA sequencing works by threading single-stranded RNA molecules through a protein pore, which can distinguish between different RNA nucleotides based on the change in electrical current caused by the passage of the nucleotides through the nanopores. In the last few years, DRS has been successfully applied to detect diverse RNA modification types, including *N6*-methyladenosine (m^6^A) (18–23), pseudouridine (Ψ) (24–26), 2′-O-methylation (Nm) (25, 27), *N7*-methylguanosine (m^7^G) (28, 29) and inosine (I) (30), and in a variety of RNA species, such as long non-coding RNAs, viral RNAs, mRNAs and tRNAs (18, 20, 31–33). Identification of modified RNA bases has been typically achieved by either: i) measuring changes in the current signal caused by the presence of modified nucleotides (20, 21, 34, 35), and/or ii) by identifying non-random basecalling ‘errors’ that occur at positions where the RNA is modified (36–39).

A central step in nanopore sequencing is the basecalling process, which involves converting the raw electrical signal data generated by the nanopore into a string of nucleotide bases that represents the RNA sequence. To convert the current intensity signal into a nucleotide sequence, the basecalling algorithm requires a basecalling model. These models are typically trained with synthetic and/or *in vivo* native RNA reads, and can predict the four canonical bases (A, C, G and U in the case of RNA). Current basecalling models, available through the *Guppy* basecalling software, are not specifically trained to predict RNA modifications, and therefore, when the model encounters a modified RNA nucleotide, it may call the wrong base (mismatch error), not call any base (deletion error) or insert a small piece of sequence (insertion error) at the modified site (18, 25, 38). Several softwares exploit this feature and use these basecalling ‘errors’ to identify RNA modifications in DRS datasets (37–41). The use of basecalling errors to detect RNA modifications is well-suited to detect certain RNA modification types, such as pseudouridine (Ψ), which causes a very strong U-to-C mismatch signature at the modified site (24–26). By contrast, other RNA modification types, such as m^6^A, yield relatively modest basecalling ‘error’ signatures (25), leading to a relatively high number of false positives and false negatives when predicting them in a transcriptome-wide fashion (18).

Basecalling errors in nanopore sequencing data can arise due to various factors, such as homopolymeric regions, RNA secondary structures or helicase mis-steps (42–44). However, a major factor affecting the basecalling ‘error’ patterns is the RNA basecalling model of choice, and more specifically, the data that was used to train this model. For example, the ‘default’ RNA basecalling model (rna_r9.4.1_70bps_hac) used in *Guppy* was trained with native mRNA sequences from diverse species, and thus contains a significant number of m^6^A modified sites in the reads that were used to train the model. Consequently, this basecalling model learnt to predict both “m^6^A’’ and “A” as “A”, leading to low frequencies in basecalling ‘errors’ when an m^6^A modification is encountered, compared to when the model encounters other RNA modification types that were not included in the training data.

Here, we hypothesized that the detection of m^6^A RNA modifications in DRS datasets would significantly improve if a RNA basecalling model trained only with unmodified bases was used. To test this, we trained novel RNA basecalling models, and then examined their performance in terms of read basecalling accuracy (i.e. sequence identity), read mappability and RNA modification detection ability. We found that an RNA basecalling model trained with fully unmodified reads, which we refer to as ‘IVT’, showed improved sensitivity for detecting m^6^A modifications. Notably, the basecalling error was mainly increased in naturally occurring m^6^A sequence contexts (DRACH, where D denotes A, G or U, R denotes A and G, and H denotes A, C or U), in agreement with our hypothesis. By contrast, other RNA modifications were detected with similar accuracy when using this model.

We also trained a second basecalling model, referred to as ‘SUP’, which we found yields 97% basecalling accuracy, compared to the 91% observed with the current RNA model. This increased accuracy was reflected in improved read mapping, especially in short RNA populations (50% increase). Moreover, we found that this model showed enhanced signal-to-noise ratio at modified bases, thus improving the detection of m1Ψ modifications in synthetic mRNA vaccines. In addition to its applicability for mRNA vaccine quality control purposes (45), this model could also be used to improve SNP detection, identification of splice sites, and transcript annotation when using DRS data.

## MATERIALS AND METHODS

### In vitro transcription

This study used synthetic RNAs known as ‘curlcakes’, which were designed to include all possible 5-mer sequences while minimizing RNA secondary structures (18), and consist of four *in-vitro* transcribed constructs: (1) Curlcake 1, 2,244 bp; (2) Curlcake 2, 2,459 bp; (3) Curlcake 3, 2,595 bp and (4) Curlcake 4, 2,709 bp. To generate ‘curlcake’ synthetic RNAs, the plasmids were digested with BamHI-HF (NEB, R3136L) and EcoRV-HF (NEB, R3195L) followed by a clean-up using phenol/chloroform/isoamyl-alcohol 25/24/1, v/v, pH = 8.05 (Sigma Aldrich, P3803). Linearized plasmids were used for *in vitro* transcription with the AmpliScribe T7-Flash Transcription Kit (Lucigen, ASF3507) using 5-Methyluridine-5’-Triphosphate (5-mUTP, Trilink, N-1024-1) instead of UTP, *N1*-Methylpseudouridine-5’-Triphosphate (m1meΨTP, Jena Bioscience, NU-890S) instead of UTP or *N4*-Acetyl-cytidine-5’-triphosphate (N4-Acetyl-CTP, Jena Bioscience, NU-988S) instead of CTP. Products were treated with Turbo DNAse (Thermo Fisher, AM2238) to remove the template plasmid and cleaned up using the RNeasy MinElute Kit (Qiagen, 74204). All constructs were polyadenylated using Escherichia coli poly(A) polymerase (NEB, M0276S) according to manufacturer’s instructions followed by a clean up using the RNA Clean & Concentrator-5 kit (Zymo Research, R1013). Concentrations were determined using the Qubit 2.0 Fluorometer. Transcript integrity, as well as polyadenylation success, were determined using an Agilent 4200 TapeStation. For previously sequenced curlcakes used in this study, including unmodified NTPs (UNM), *N6*-methyladenosine triphosphate (m^6^ATP), 5-methylcytosine triphosphate (m^5^CTP), 5-hydroxymethylcytosine triphosphate (hm^5^CTP) and pseudouridine triphosphate (ΨTP), publicly available raw fast5 files were used as input for the analysis (18, 25) (see **Table S2)**.

### Direct RNA sequencing of *in vitro* transcribed RNA

The RNA libraries for direct RNA sequencing of ac^4^C, m^5^U, m^1^Ψ-containing curlcake constructs and UNM-S (which included Bacillus subtilis guanine riboswitch, B. subtilis lysine riboswitch and Tetrahymena ribozyme) were prepared following the ONT Direct RNA Sequencing protocol version DRS_9080_v2_revR_14Aug2019-minion. Briefly, 800 ng of poly(A)-tailed RNA (200 ng per curlcake construct and 250 ng per riboswitch) was ligated to ONT RT Adaptor (RTA) using concentrated T4 DNA Ligase (NEB, M0202T). The optional reverse transcription step was performed using SuperScript III (Thermo Fisher Scientific, 18080044) in the case of ac^4^C, m^5^U-containing curlcake constructs and UNM-S, and Superscript IV (Thermo Fisher Scientific, 18090010) in the case of m^1^Ψ. The products were purified using 1.8x Agencourt RNAClean XP beads (Thermo Fisher Scientific, NC0068576) and washed with 70% freshly prepared ethanol. RNA Adapter (RMX) was ligated onto the RNA:DNA hybrid, and the mix was purified using 1× Agencourt RNAClean XP beads, followed by washing with wash buffer (WSB) twice. The sample was then eluted in elution buffer (ELB) and mixed with RNA running buffer before loading onto a primed R9.4.1 flowcell. The samples were run on a MinION sequencer for 24-72h. The sequencing runs were performed on independent days and using a different flowcell for each sample.

### Training a high-accuracy RNA basecallers (SUP model)

We trained a new RNA basecalling model (SUP) using bonito v0.5.1 (https://github.com/nanoporetech/bonito). SUP model is based on the latest CTC-CRF (bonito) DNA model architecture, which achieves 96% accuracy for DNA basecalling with chemistry R9.4.1. DNA CTC-CRF models use a window of 19 and stride of 5. For our RNA models, we used a window of 31 and stride of 10 (similar to RNA flip-flop models).

Bonito model training requires chunks of fixed length (10,000 samples) and underlying sequences. We kept only chunks that corresponded to 30-270 bases. Chunks from each read were generated by stepping every 50 bases in the sequence space. Training was performed using bonito by keeping 3% of randomly selected chunks for validation. The SUP model was trained using publicly available human datasets containing triplicates of wildtype and *Mettl3* KO cells (21). We kept the 5 longest reads per transcript and run in order to balance highly and lowly expressed transcripts. In total, 158,413 reads were used for training. The training was stopped after 5 epochs. We selected the model with the lowest validation error as the final one from each training, corresponding to epoch 5 for the SUP model. The resulting model was calibrated using the *calibrate_qscores_byread.py* script from taiyaki, and reached 97% median accuracy on native human RNA (**Fig 1B**).

**Figure 1.**
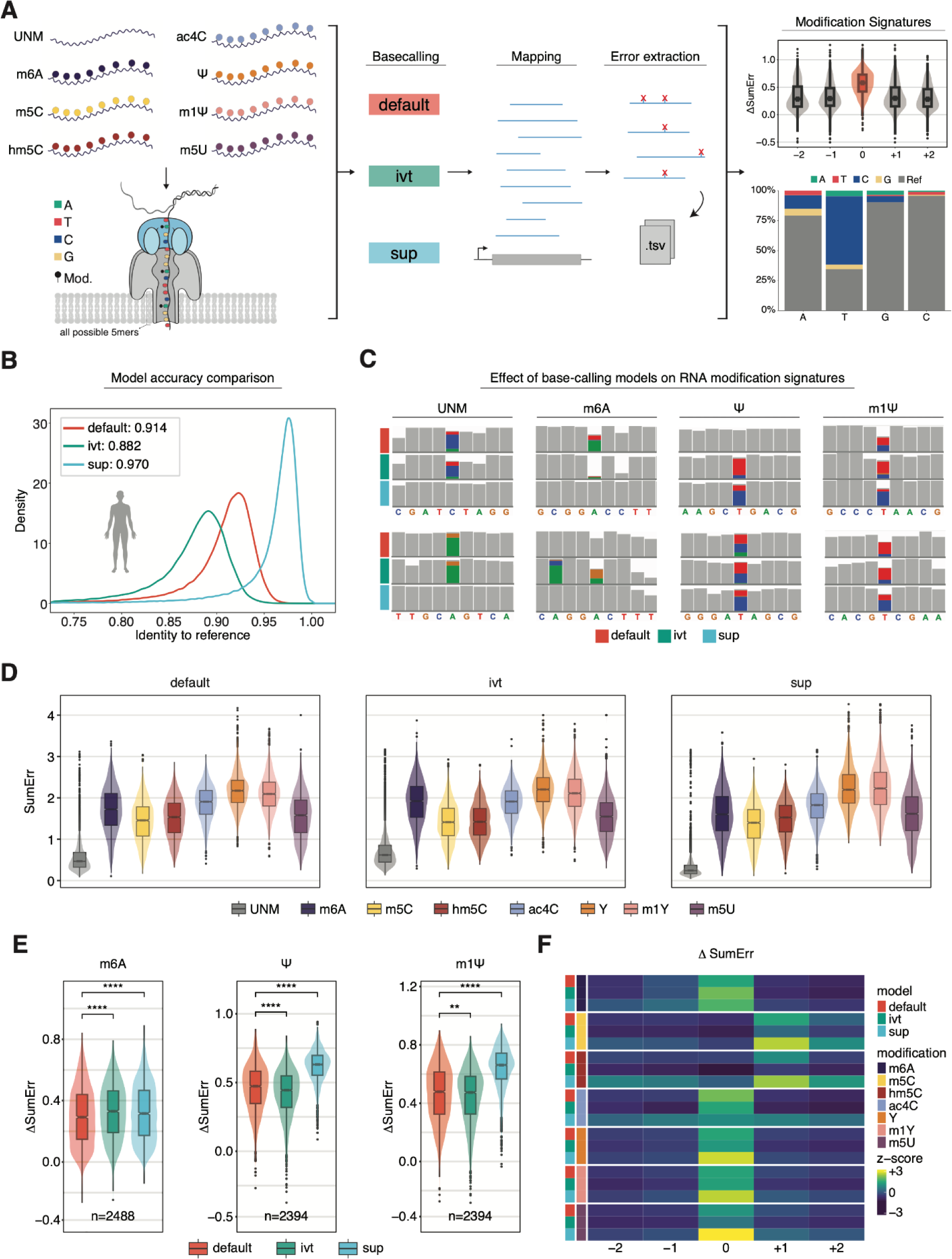
Benchmarking RNA basecalling models and their effect on basecalling accuracy and RNA modification detection. **(A)** Schematic overview of the *in vitro* ‘curlcakes’ used in this work, depicting the 7 different modification types that were sequenced (m^6^A, m^5^C, hm^5^C, ac^4^C, Ψ, m^1^Ψ and m^5^U) and the three RNA basecalling models benchmarked (default, IVT, and SUP). To assess the performance of tested RNA basecalling models, error signatures (mismatch-, deletion-, and insertion frequency) were used. **(B)** Comparison of basecalling accuracies of the distinct RNA basecalling models used in this work. The curve represents the density of the per-read identities obtained from HEK293T wildtype cells (rep1) mapped to the human transcriptome (see *Methods*). **(C)** IGV snapshots showing the effect of using either default, IVT or SUP models on unmodified (UNM) or modified (m^6^A, Ψ and m^1^Ψ) ‘curlcake’ sequences. Positions at which the mismatch frequency exceeds 0.2 are colored, while deletions are visualized as drop in coverage. **(D)** Comparison of the summed errors (mismatch, deletion, and, insertion frequency) for all 5-mers with a central modified base for the three tested basecalling models (default, IVT and SUP). **(E)** Boxplots showing the delta summed error of modified (position 0) 5-mers for selected modifications (m^6^A, Ψ and m^1^Ψ). See Figure S2B for equivalent plots of the remaining RNA modifications tested (m^5^C, hm^5^C, ac^4^C and m^5^U). Statistical analysis was performed using a two-sided non-parametric Wilcoxon test with ‘default’ as the reference group. Results were corrected for multiple-hypothesis testing using the Benjamini-Hochberg procedure to obtain adjusted p-values (ns: p > 0.05, *: p <= 0.05, **: p <= 0.01, ***: p <= 0.001, ****: p <= 0.0001). The sample size reported by *n* represents the number of 5-mers contributing to each boxplot per panel. For Figures 1D and 1E the box is limited by the lower quartile Q1 (bottom) and upper quartile Q3 (top). Whiskers are defined as 1.5 * IQR with outliers represented as individual dots. **(F)** Comparison of the median delta summed error (= SumErr_MOD_ - SumErr_UNM_) for centrally modified (position 0) 5-mers. A z-score normalization per modification (see *Methods*) was performed to highlight the effect that different basecalling models have on the obtained delta summed error. Unscaled results are reported in Figure S2B.

### Training an RNA basecaller using *in vitro* sequences (IVT model)

The IVT model was trained using publicly available IVT direct RNA data obtained from CEPH1463 cells (UCSC_Run1_IVT_RNA). RNA model training procedure was performed for 50,000 iterations with following taiyaki hyperparameters: --size 256 --stride 10 --winlen 31 --chunk_len 4000 --batch_size 50. We selected up to 100 chunks from every read using a step of 25 bases. IVT model was trained using taiyaki v4.1.0 (https://github.com/nanoporetech/taiyaki).

### Demultiplexing direct RNA sequencing

To evaluate the effect of differing m^6^A-stoichiometries on the sensitivity of each basecalling model we used publicly available, multiplexed curlcake data (PRJEB61874) containing a range of stoichiometries (barcode1 = 12.5%, barcode 4 = 25%, barcode 3 = 50% and barcode 2 = 75%, m^6^ATP). Demultiplexing of the barcoded direct RNA sequencing libraries was performed using DeePlexiCon (46) keeping reads with demultiplexing confidence scores above 0.80 (`-f multi -m resnet20-final.h5 -s 0.8`) .

### Basecalling and mapping of direct RNA sequencing datasets

Publicly available datasets for *H. sapiens (21)*, *M. musculus (30)*, *X. laevis (30)*, *S. cerevisiae* (18), *A. thaliana* (41) and the synthetic eGFP mRNA vaccine (45) were downloaded from their respective data repositories reported in **Table S2.** Raw fast5 files were basecalled and mapped using the MasterOfPores pipeline (version 2.0) (47). For basecalling, *Guppy* (version 6.0.6) was used with either the default, IVT or SUP model (**Table S4**). For highly modified curlcake runs and the eGFP mRNA vaccine, basecalled fastq files were aligned to the curlcake reference using graphmap (48) (version 0.5.2) with *‘sensitive*’ settings (25) (`--rebuild-index -v 1 --double-index --mapq -1 -x sensitive -z 1 -K fastq --min-read-len 0 -A 7 -k 5`). For the *in vivo* m6A-analysis basecalled fastq files were aligned to the soft-masked human genome (Homo_sapiens.GRCh38.dna_sm.primary_assembly.fa) using minimap2 (v2.17) with *‘default’* settings (`-uf -ax splice -k14`) . Uniquely mapped reads were reported with Samtools (49) (version 1.15.1) using `samtools view -c -F 260` (**Table S4**).

For the cross-species comparison reads were mapped to the respective reference transcriptome (**Table S4)** using minimap2 (v2.17) with *‘transcriptome’* settings (`-ax map-ont -k14`). Aligned reads were first filtered for uniquely mapped reads with a baseQ > 15 using Samtools (49)(`samtools view -q 15 -F 3840 <in.bam> > <out.bam>`). To account for the heterogeneous expression profiles of each transcriptome and to prevent biasing our analysis towards highly expressed genes we sampled the longest reads per transcript keeping at most 10 reads per transcript (`bam2select.py`). Per read identities were calculated using a custom python script (`identity_density_v2.py`). All scripts required to reproduce results in this manuscript are provided on the project’s github page (https://github.com/novoalab/basecalling_models). Silhouettes of model species were downloaded from PhyloPic (https://www.phylopic.org/).

### Comparative analysis of basecalled features in curlcakes

To qualitatively assess the difference in basecalling errors introduced by default, IVT and SUP upon RNA modifications, the sorted and indexed bam files were visualized using the Integrative Genomics Viewer (IGV, version 2.13.1) (50). Using EpinanoRMS (25) (version 1.1) basecalled features (mismatch frequency, deletion frequency and insertion frequency) were extracted from the alignment files on a per-position basis. To obtain per 5-mer information this output was converted using `epinano_to_kmer.R` script which applies a sliding window of a given size to compile the per k-mer information. These results were further processed to calculate the summed error (Σ(mismatch frequency, deletion frequency, insertion frequency)) and delta summed error (SumErr_MOD_ - SumErr_UNM_) on a per-position basis and visualized using i*n-house* R scripts. To calculate the mismatch directionality the number of reads corresponding to the correct reference base for that position was divided by the total number of reads obtained for that position. All scripts required to reproduce these results are available under the project’s GitHub repository (https://github.com/novoalab/basecalling_models).

### Detection of m^6^A-modified sites in HEK293T using comparative error analysis

To determine whether alternative basecalling models can improve the detection of m^6^A *in vivo,* the alignment files of individual replicates for wildtype and *Mettl3^-/-^* were merged to obtain greater coverage. To reduce computational load, only protein-coding genes were considered in the comparison. For this reason, lines in which the gene biotype corresponds to protein coding were extracted from the genome annotation file and converted to bed format using `bedops gtf2bed` (version 2.4.41) (51). The resulting bed file was split into individual chromosomes which would serve as regions for testing in the pairwise comparison step. To perform pairwise comparisons, we ran eligos2 (v2.1.0) (38) using the singularity image provided on the software’s GitLab repository (https://gitlab.com/piroonj/eligos2) with a p-value cut-off of 1e-5 an odds Ratio of 2.5 and each of the individual chromosome files as regions for testing. Next, results obtained from individual chromosomes were merged using an in-house shell script and filtered to remove non-A bases as well as homopolymers to identify putative m^6^A sites (`eligos2 filter -sb A --homopolymer`).

### Metagene analysis and motif identification

To visualize the transcriptome-wide distribution of putative m^6^A sites identified with distinct basecalling models we used the GuitaR package (version 2.12.0) (52). For *de novo* motif discovery, sites obtained from eligos2 were converted into fasta format which contained 6bp up and downstream of the modified site using the `table2fa_eligos.mergeextend.sh` script. Subsequently, those files were used for *de novo* motif prediction using BaMM motif (53) (version 1.4.0) using default settings and a z-score threshold of 5.

### Comparison of identified m^6^A-sites against m^6^ACE-seq, GLORI-seq and miCLIP-seq

To identify high confidence m^6^A-sites we intersected results obtained by each of the three models (default, IVT and SUP) with data obtained from orthogonal methods. These datasets included m^6^ACE-seq data that was matched with the nanopore results, as well as GLORI-seq and miCLIP-seq results obtained from the same cell line (21, 54, 55). These datasets were filtered to only retain sites for which we had sufficient coverage in the nanopore runs (≥ 20x) within protein coding genes (https://github.com/novoalab/basecalling_models). Overlaps between datasets were visualized using the ggvenn R package (version 0.1.9).

For stoichiometric analysis, sites overlapping with GLORI-seq were used as this dataset provides per-nucleotide resolution information as well as an estimated stoichiometry for each identified m^6^A-site (54). To this end, results obtained by GLORI-seq were first binned based on their reported stoichiometry into sites that were either considered HIGH (≥0.75 modified), MED-HI (≥0.50 and <0.75 modified), MED-LO (≥0.25 and <0.50 modified) or LOW (≤0.25 modified). Only those sites were retained in the final set in which both GLORI-seq replicates were reported within the same bin. This resulted in a total of 47722 high-confidence sites split between HIGH (n=8900), MED-HI (n=7947), MED-LO (n=12697) and LOW (=18178).

### Processing of eGFP mRNA vaccine data and comparative analysis

Aligned reads were filtered to obtain the 20.000 longest reads and matched between default and SUP (`bam2select.py`). Mismatch-, deletion- and insertion-frequencies were calculated and extracted on a per nucleotide basis using EpinanoRMS (18).The per base frequencies were calculated by dividing the number of each of the four canonical nucleobases (A, C, G or T) for each position by the number of total observations for that position. To determine differences in sensitivity between the two models, we extracted the 20 most lowly U>C modified sites in default and compared them to the same positions in SUP. All scripts available at https://github.com/novoalab/basecalling_models.

## RESULTS

### RNA basecalling model choice affects both per-read accuracy and RNA modification ‘errors’

Currently, detection of RNA modifications in DRS datasets is achieved via two different approaches: i) identification of current signal alterations, or ii) increased basecalling ‘errors’ (36). The latter approach has been exploited by diverse softwares and algorithms, such as EpiNano (18), DiffErr (41), ELIGOS2 (38), DRUMMER (56) and JACUSA2 (39), among others. All of these methods are affected by the choice of RNA basecalling model used, which is responsible for calling these ‘errors’ when it encounters a modification.

We hypothesized that the use of alternative basecalling models might enhance the detection of RNA modifications by increasing the ‘error’ signatures observed in the basecalled data. To this end, we trained novel RNA basecalling models using two different approaches (**Fig S1A**), to assess their effect on both basecalling accuracy as well as on basecalling ‘error’ patterns, which can be used as proxy for RNA modifications (**Fig 1A**). The first approach consisted in training a model using fully unmodified RNA sequences generated via *in vitro* transcription (‘IVT’ model), which we reasoned should lead to increased ‘error’ signatures at modified bases present in mRNAs, compared to the default basecalling model. The second approach consisted in generating a ‘super-accuracy’ RNA basecalling model (‘SUP’ model) (see **Table S1**), which we reasoned would show increased basecalling accuracy (**Fig S1B**), and consequently, improved detection of RNA modifications due to a increased signal-to-noise ratio.

Once the models were trained, we first compared the performance of each model on human HEK293T cells, finding that the SUP model was the most accurate out of the three models tested, reaching a median read accuracy of 97% on human cells (**Fig 1B**), while the default and IVT models only showed 91% and 88% accuracy, respectively. We then examined how basecalling ‘error’ signatures produced by m^6^A, m^5^C, hm^5^C, ac^4^C, Ψ, m^1^Ψ, and m^5^U in synthetic ‘curlcakes’ (18) would vary depending on the RNA basecalling model choice (default, IVT or SUP) (**Fig 1C**, see also **Fig S1C)**. Of note, ‘curlcakes’ are synthetic constructs that contain all possible 5-mers, thus allowing to study the effect in all possible sequence contexts that affect the basecalling process. To this end, we employed per-position mismatch, deletion and insertion errors, which we consider jointly as ‘summed errors’ (SumErr) (see *Methods*). Our analyses revealed that SumErr values in unmodified reads were significantly decreased when basecalled with the SUP model (**Fig 1D, Fig S2A**), in agreement with the increased accuracy of the model.

We then assessed the ability of each model to detect RNA modifications in the form of basecalling errors. For this purpose, we computed the delta summed error (ΔSumErr = SumErr_MOD_-Sumerr_UNM_) for each position, model and modification type, which corrects for the different background error rates of each model on a per-5-mer basis. Our results showed that, for most modifications examined –and especially for m^5^U, Ψ and m^1^Ψ–, the ΔSumErr was significantly increased when using the SUP model, relative to default (**Fig 1E**, see also **Fig S2B**), indicating that the signal-to-noise ratio was increased, and thus confirming the hypothesis that the use of alternative basecalling models can improve our ability to detect RNA modifications in DRS data. By contrast, the use of the IVT model did not increase the ΔSumErr for most RNA modifications, with the exception of m^6^A basecalling errors, which were modestly –yet significantly– increased when using the IVT model, compared to default (**Fig 1E**), in agreement with our hypothesis.

In addition, we noticed that all RNA modifications, regardless of the basecalling model, showed increased basecalling errors at the modified site (position 0), with the exception of m^5^C and hm^5^C, which showed increased errors at position +1, in agreement with previous reports (25). Moreover, we observed that for m^6^A the increase in “error” using the IVT model compared to the default model was mainly due to high deletion signatures at the modified site (**Fig S2C-D**). On the other hand, the increased signal in SUP model to detect Ψ and m^1^Ψ modifications was caused by an increase in the global U>C mismatch frequency in SUP, compared to default (**Fig S2C-D,** *see also* **Fig S3**). Taken together, these results show that SUP and IVT basecalling models achieve a stronger signal-to-noise ratio either via changing the error signature (IVT model for m^6^A) or improving previously observed patterns (SUP model for Ψ and m^1^Ψ).

### IVT model outperforms other models in detecting m^6^A modifications in DRACH contexts

*N6*-methyladenosine is the most prevalent epitranscriptomic mark in mammalian mRNA molecules, with an average of 1-3 sites per transcript (7, 8). Thus, the use of native RNA molecules as training dataset for a basecaller (as is the case for both default and SUP models, see **Fig S1**) will lead to basecalling models learning to interpret both “m^6^A” and “A” as “A”, since the training data contains large amounts of m^6^A-modified residues while its output is limited to the four canonical nucleobases (A, C, G and U). For this reason, we speculated that the relative basecalling ‘error’ rates in our IVT model should be particularly improved in those sites that naturally contain m^6^A (DRACH, where D denotes A, G or U, R denotes A and G, and H denotes A, C or U) and are therefore masked in ‘default’.

To confirm this, we independently examined the performance of the 3 models in DRACH and non-DRACH motifs. Indeed, we observed that the modest increase in IVT model performance at m^6^A-containing sites (**Fig 1E**) was significantly stronger when only DRACH sites were taken into account (**Fig 2A**), in agreement with our hypothesis. A detailed examination of which individual 5-mers were contributing to this phenomenon revealed that certain DRACH 5-mers showed no or only mild improvement while others were strongly enhanced in their signal-to-noise ratio (**Fig 2B, Fig S4A**). Notably, the largest improvement in the signal difference was observed for GGACT (**Fig 2B**, see also **Fig S4B**) which is the most prevalent m^6^A-motif observed in mammalian species (57), suggesting that mammalian native RNA was most likely used to train the ‘default’ basecalling model.

**Figure 2.**
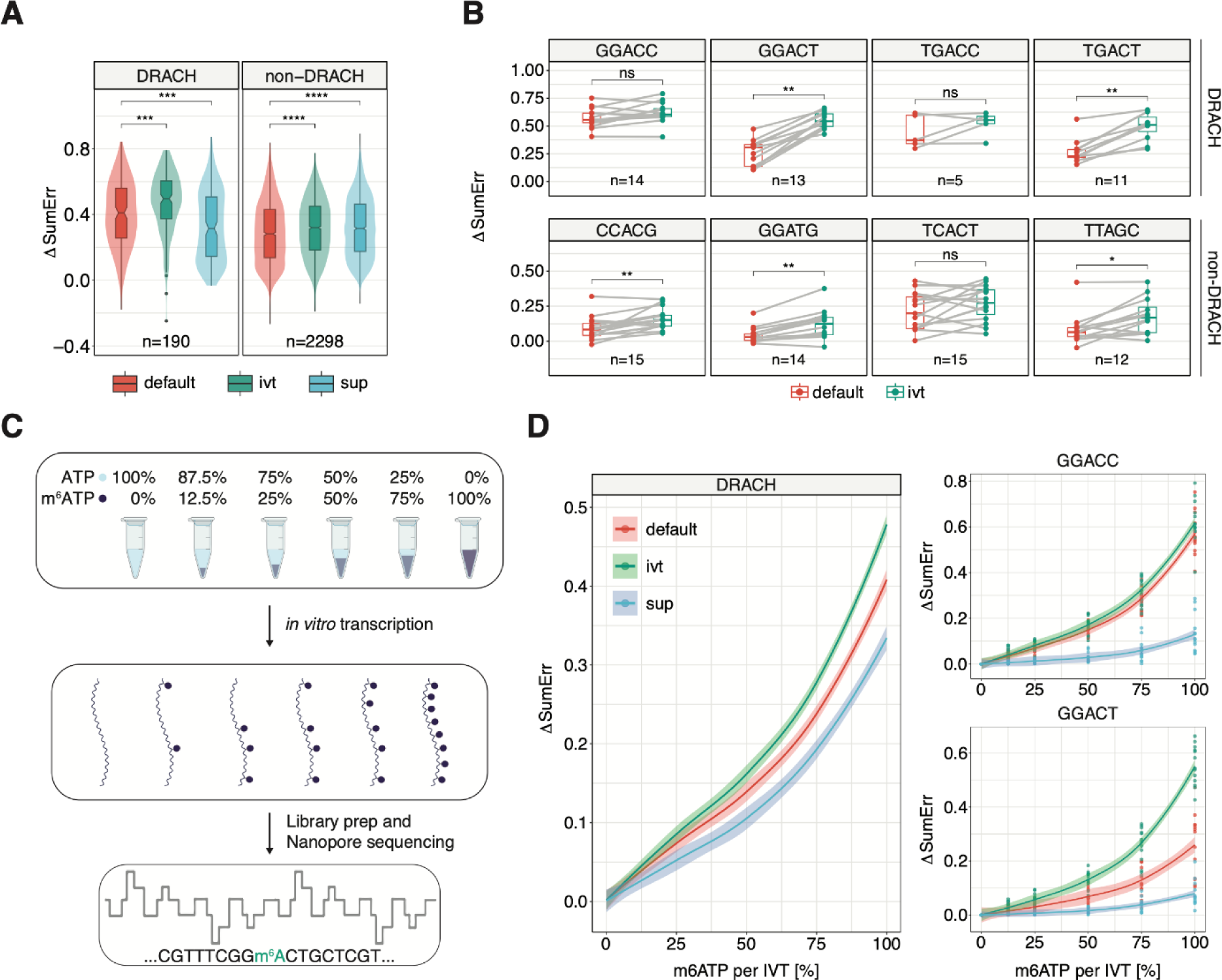
The effect of custom basecalling models on biologically relevant m^6^A motifs and at different m^6^A stoichiometries. **(A)** Comparison of the summed errors (=Σ mismatch-, deletion-, and, insertion frequency) at the central modified position for 5-mers that fall within the DRACH motif (D=A, G or U; R=G or A; H=A, C or U) and others (non-DRACH). Statistical analysis was performed using a two-sided non-parametric Wilcoxon test with ‘default’ as the reference group. **(B)** Comparison of the delta summed errors (= SumErr_MOD_ - SumErr_UNM_) for individual motifs that are DRACH and non-DRACH. Each dot represents a single observation of that 5-mer within the ‘curlcake’ sequences. To compare the relationship between individual 5-mers a paired two-sided Wilcoxon test was performed. **(C)** Schematic representation of the experimental approach to obtain ‘curlcakes’ at different m^6^A stoichiometries. ATP and m^6^ATP were mixed at different ratios and used for the *in-vitro* transcription of ‘curlcakes’ with the remaining NTPs, followed by poly-A-tailing, library preparation, and direct RNA sequencing. **(D)** Comparison of error-signature amplitude at different modification stoichiometries between 5-mers that fall within the DRACH motif. The corresponding plot for non-DRACH motifs is reported in Figure S5B. Lines were fitted using locally estimated scatterplot smoothing (loess) and lightly shaded areas correspond to the 95% confidence interval. For Figures 3A-B the box is limited by the lower quartile Q1 (bottom) and upper quartile Q3 (top). Whiskers are defined as 1.5 * IQR with outliers represented as individual dots. All statistical tests were corrected for multiple hypothesis testing using Benjamini-Hochberg correction and adjusted p-values are displayed (ns: p > 0.05, *: p <= 0.05, **: p <= 0.01).

Finally, we examined the sensitivity of the different basecalling models with varying modification stoichiometries. To this end, we generated ‘curlcak’ *in vitro* transcripts at varying modification levels by mixing ATP and m^6^ATP at different ratios (**Fig 2C**, see also *Methods*). Our results showed that even at the lowest modification level (12.5%), the IVT model was able to generate a significantly increased error signature at DRACH-bearing 5-mers compared to the default model, which was especially evident in the GGACT context (**Fig 3D**, see also **Fig S5A-C).**

**Figure 3.**
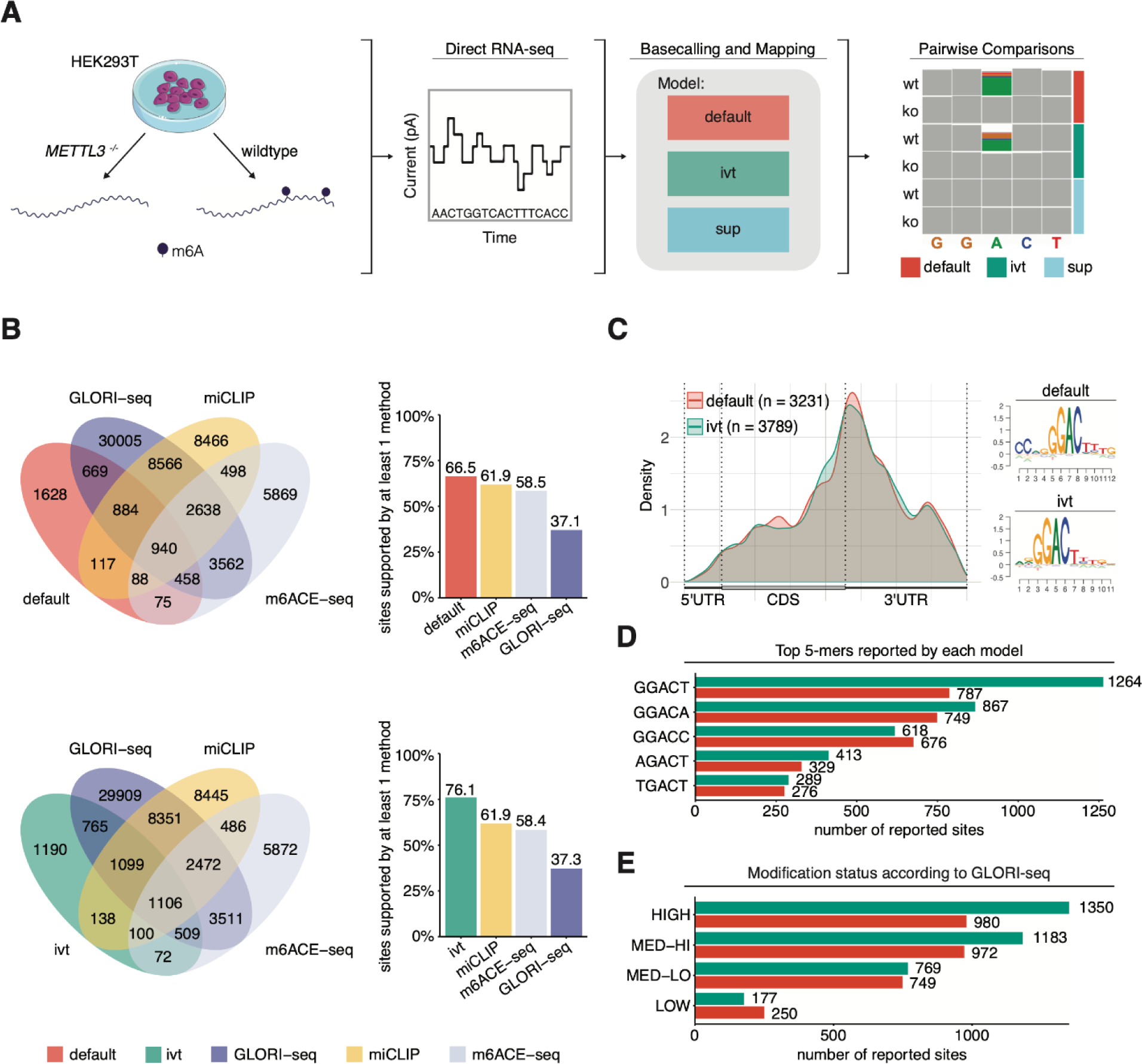
Detection of m^6^A-modified sites from HEK293T cells using a pairwise comparison approach and validation of identified sites with orthogonal methods. **(A)** Schematic representation of the workflow to obtain m^6^A-modified sites using direct RNA-sequencing. Samples from HEK293T wildtype and *METTL3 ^-/-^*knockout cells were sequenced using direct RNA-sequencing. Raw fast5 files were basecalled using either of the three models tested (default, IVT and SUP) and mapped to the human reference genome (hg38). To obtain m^6^A-modified sites. pairwise comparisons within each model were performed using eligos2 (38). **(B)** Venn diagrams showing the overlap in absolute numbers between nanopore used with either default or IVT model and m6ACE-seq, miCLIP and GLORI-seq (21, 54). Barplots show the relative amount of sites supported **(C)** Metagene plot of sites identified by either default or IVT model show the known pattern of m^6^A distribution with a strong enrichment around the stop codon. Corresponding motifs identified *de novo* using BaMM. **(D)** Barplot showing the top 5-mers identified by both models. **(E)** Sites overlapping with GLORI-seq were binned according to their modification stoichiometry reported by GLORI-seq (HIGH: ≥0.75 modified; MED-HI: ≥0.50 and <0.75 modified; MED-LO: ≥0.25 and <0.50 modified; LOW: ≤0.25 modified).

### Improved *in vivo* detection of m^6^A-modified sites in human samples using IVT model

Next we examined whether the IVT model would improve the detection of modified sites *in vivo* using third-party tools for the detection of differentially modified m^6^A sites. Therefore we used publicly available nanopore DRS data from HEK293T wildtype and *Mettl3*^-/-^ cells (21, 54), basecalled the data using our three basecalling models, mapped the reads to the reference genome, and finally used ELIGOS2 (38), a tool developed to identify RNA modifications through pairwise comparison of error profiles, to detect m^6^A sites (**Fig 3A**). Comparison of control and *Mettl3^-/-^*samples using ELIGOS2 revealed a similar amount of sites detected when basecalling the data using default (4,859 m^6^A sites) and IVT model (4,979 m^6^A sites) (**Table S3**), showing that the use of the IVT model modestly increased the number of predicted m^6^A modified sites.

To further examine this point, we compared the set of ELIGOS2-predicted m^6^A sites to a list of m^6^A sites predicted in HEK293T cells using orthogonal methodologies (m6ACE-seq (21), GLORI-seq (54) and miCLIP (58)). This analysis revealed that 67.% of the sites reported by ELIGOS2 on reads basecalled with default model were supported by at least one orthogonal method, whereas 76% of the sites reported by the IVT model had orthogonal support (**Fig 3B**, see also **Fig S6A-C**). Sites that were supported by at least one orthogonal method demonstrated the typical m^6^A distribution along the transcript, and motif enrichment of the identified m^6^A sites matched the mammalian m^6^A motif, suggesting that these sites are likely true m^6^A sites (**Fig 3C, Fig S6D**). Furthermore, we noted that m^6^A sites predicted with IVT model captured a higher proportion of GGACT sites compared with those predicted when basecalling with default model (**Fig 3D**, see also **S6E**). We then further examined whether the predicted m^6^A sites corresponded to high or low stoichiometry m^6^A sites. To this end, we binned the sites based on the modification stoichiometry values predicted by GLORI-seq, finding that the IVT model predicted a larger proportion of high stoichiometry sites, suggesting that sites predicted with IVT-basecalled data contain a higher proportion of true positives, compared to default (**Fig 3E**, see also **Fig S6E**). These results show that ELIGOS2-predicted m^6^A sites on IVT basecalled datasets are more accurate at predicting m^6^A sites compared to ELIGOS2-predicted m^6^A sites on default basecalled datasets. In addition, the IVT model was capable of detecting unique m^6^A sites that were neither reported by any of the orthogonal methods nor the other two basecallers tested, although these sites are supported by the most comprehensive m^6^A database available (59), suggesting that IVT can identify unique sites (**Fig S6F**). By contrast, the same comparison performed with SUP reported only a very small number of m^6^A sites (97), which lacked overlap with the ground-truth datasets (**Fig S6B-C**), suggesting that either SUP is not suited for the detection of m^6^A using the pairwise-comparison of error-profiles, or that ELIGOS2 software –or the default parameters used– are not suited for models that do not show strong ‘error’ patterns.

### SUP basecalling model shows 97% read accuracy and up to 50% increase in mapped reads

Nanopore sequencing shows relatively high error rates compared to other sequencing platforms, especially in the case of direct RNA sequencing, with ∼12% per-read error rates (15). This high background limits the technology’s applicability in several areas such as SNP detection, transcript annotation and RNA modification detection. In addition to changes in pore chemistry that can mitigate sequencing errors, the introduction of basecalling models with super-high accuracy represents an alternative approach to enhancing the technology’s accuracy.

To this end, we examined the per-read accuracy of the different models trained, across a wide range of model species, using previously published DRS datasets (21, 25, 30, 41), and found a significant increase in median read accuracy of SUP compared to the default model, with the SUP model reaching a median accuracy of 97% in human samples (**Fig 4A**, see also **Fig S7A**).

**Figure 4.**
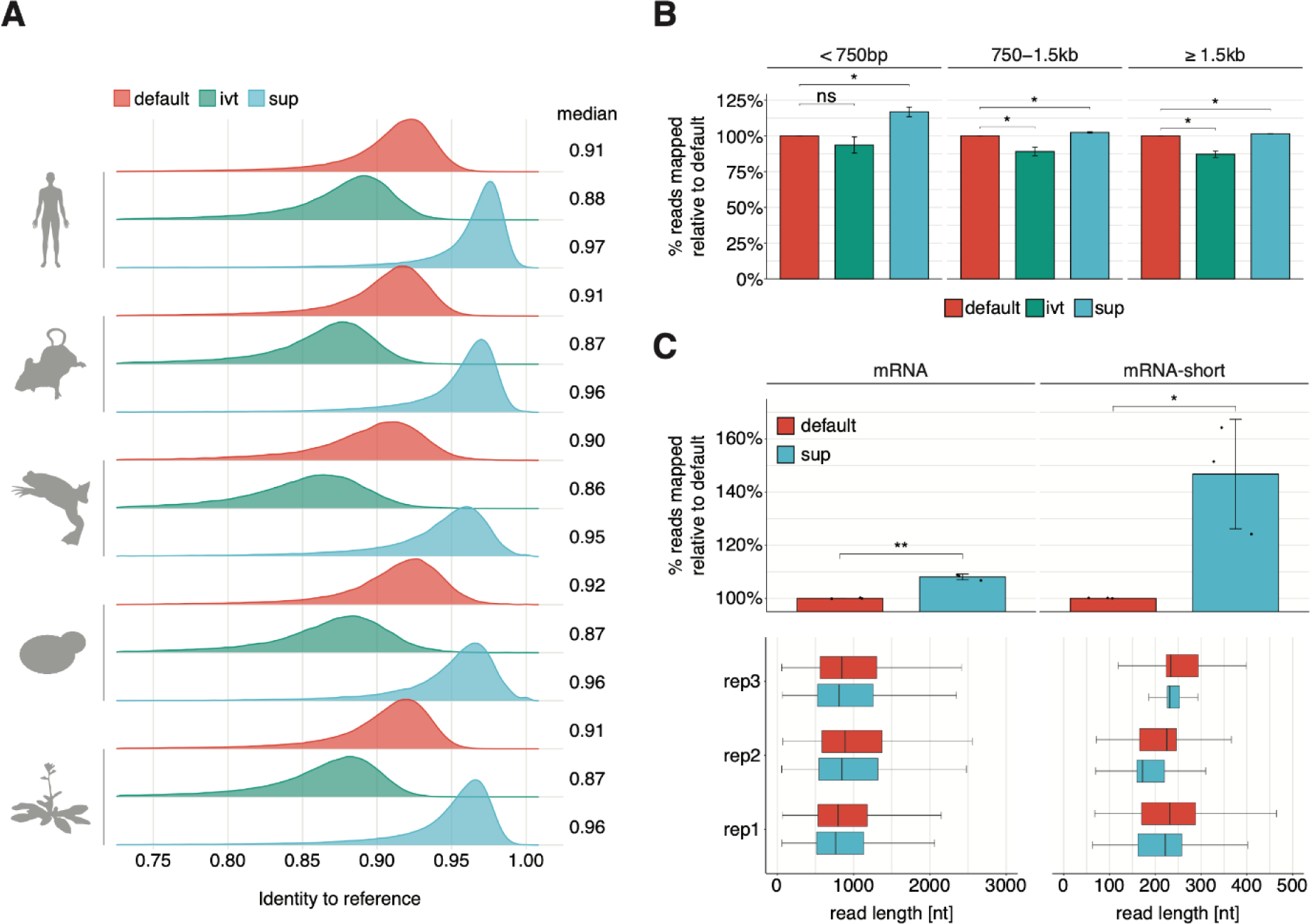
SUP is a super high-accuracy model that leads to improved read accuracy and mapping. **(A)** Cross species comparison of read accuracies obtained from each model. Reads were aligned to the respective reference transcriptome (see **Table S4**). To account for vastly different expression levels that would bias the analysis towards strongly expressed genes we filtered for the longest reads of each transcript keeping at most 10 reads per transcript. **(B)** Comparison of mapped reads obtained from HEK293T wildtype cells (n = 3), across three bins of different read lengths. Reads were mapped to the human reference transcriptome. Error bars indicate +/− 1 sd. Statistical significance was determined using two-sided t-test, corrected for multiple hypothesis testing using the Benjamini Hochberg procedure (ns: p >= 0.05, *: p < 0.05). **(C)** Effect of basecalling model on the proportion of mapped reads. Data shown corresponds to publicly available DRS data on polyA-selected HEK293T samples (left) and an mRNA-short library (mix of three synthetic sequences that were *in-vitro* polyadenylated, see also *Methods*). Statistical significance was determined using one-sided t-test, and results were corrected for multiple hypothesis testing using the Benjamini Hochberg procedure (ns: p >= 0.05, *: p < 0.05, **: p < 0.01). Boxplots depict the underlying size distribution of the sequenced libraries. The box is limited by the lower quartile Q1 (bottom) and upper quartile Q3 (top). Whiskers are defined as 1.5 * IQR with outliers being removed.

We then examined whether the increase in read accuracy would be reflected in the form of an improvement in read mappability. We hypothesized that this should particularly impact shorter reads, as errors in those will be penalized more by most mapping algorithms, and may lead to reads remaining unmapped. To test this, we examined the relative proportion of mapped reads compared to default, binning the reads by their read length. Notably, we found a consistent and significant increase in mappability for all bins tested when reads were basecalled with the SUP model, with the difference with default being the largest in the shortest bin (< 750bp category; relative increase compared to default = 117%) **(Fig 4B**, see also **Fig S7B)**. We then examined whether this phenomenon would become even more evident in very short RNAs (mRNA-short, median length ≅ 200nt), which are often observed in highly fragmented samples, such as blood plasma or sperm (60, 61). This analysis revealed that the increase in mappability was ∼5X larger in the short mRNA library (relative increase compared to default = 146%) **(Fig 4C**, *see also* **Fig S7C**).

### Enhanced U>C mismatch signature and reduced background error in a vaccine model using a SUP model

Recently, the need for an improved quality control pipeline to characterize mRNA vaccine production has been pointed out (45). In this regard, nanopore direct RNA sequencing offers a unique platform as it can simultaneously provide information about sequence identity, RNA integrity, poly-A-tail length and modification status. Here, we wondered whether the use of the SUP model, which produced very high U>C conversions at m^1^Ψ-modified sites (55 %) in the ‘curlcake’ sequences (**Fig S8**) and increased basecalling accuracies (**Fig 4A**), would be useful to characterize m^1^Ψ-containing mRNA vaccines, such as those recently used against COVID-19 (62). More specifically, we wondered whether the use of SUP could reduce the background error (improve basecalling accuracy and mappability) while at the same time improving the detection of modified nucleotides in an mRNA vaccine. To this end, we re-examined publicly available m^1^Ψ-modified and unmodified matched DRS data, which consisted of a synthetic mRNA vaccine that contained an alpha-globin 5’ UTR, an enhanced green fluorescent protein coding sequence and an AES-mtRNR1 3’ UTR (45). Our results revealed that all synthetic sequences showed reduced background error when basecalled with our SUP model, both in the m^1^Ψ-modified and unmodified variants (**Fig 5A**, see also **Fig S9A**), while showing a significant increase in the mismatch error (**Fig 5B** *see also* **FigS9B**), typically in the form of an increased U>C error (**Fig S9C**). Indeed, out of a total of 170 m^1^Ψ-modified sites, we observed an increased U>C error for 151 sites (≈ 80%) in SUP over default (**Fig 5C**). More importantly, for sites that previously showed a very poor signature in default we observed a global improvement in U>C conversion suggesting that our SUP model is more sensitive towards m^1^Ψ (**Fig 5D**, *see also* **FigS9D**). Overall, our results suggest that SUP model can be a preferable basecalling model for performing quality control on mRNA vaccines and testing the incorporation of modified nucleotides, due to its reduced background error and increased signal-to-noise ratio at m^1^Ψ-modified sites.

**Figure 5.**
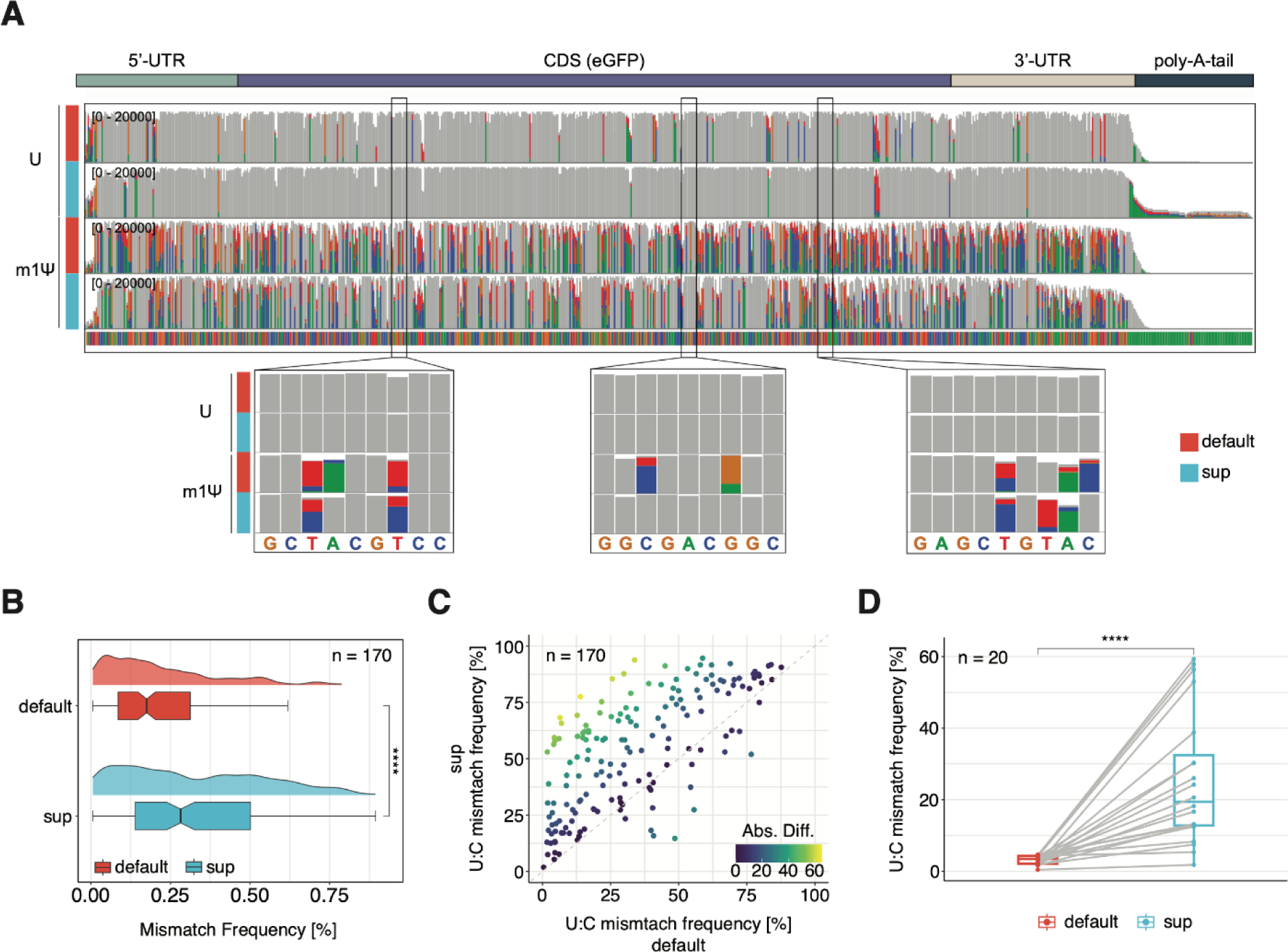
SUP improves the identification of m1Ψ-modified residues in synthetic mRNA vaccines. **(A)** IGV screenshots of the synthetic eGFP vaccine in unmodified (U) and modified (m1Ψ) molecules, basecalled with either default (red) or sup (cyan) basecalling models. The three zoomed regions showcase the increased U>C conversion rate at m1Ψ sites as well as the reduced background error produced by the sup model. The 20,000 longest, uniquely mapped reads were selected for this analysis. Positions at which the mismatch frequency exceeds 0.2 are colored while those with a mismatch frequency below 0.2 are shown in gray. **(B)** Comparison of mismatch frequency across all m1Ψ sites (n = 170) found in the synthetic eGPF vaccine. To test for statistical significance the non-parametric Wilcoxon-test was used and values corrected for multiple hypothesis testing using the Benjamini-Hochberg procedure.(ns: p > 0.05, *: p <= 0.05, **: p <= 0.01, ***: p <= 0.001, ****: p <= 0.0001). **(C)** Comparison of U>C mismatch frequency for all 170 m1Ψ sites between sup and default. Points located in the upper half of the dotted line represent sites with higher U>C frequency in sup (n = 151) while points below the dotted line represent sites with higher U>C frequency in default. (n = 19). Points are colored by the absolute difference between sup and default. **(D)** Comparison of U>C mismatch frequency across the most lowly modified sites in default (median = 3.45) demonstrates the improved sensitivity of sup (median = 19.43) in detecting m1Ψ. To compare the relationship between individual positions a paired two-sided Wilcoxon test was performed. For Figures 5B and 5D the box is limited by the lower quartile Q1 (bottom) and upper quartile Q3 (top). Whiskers are defined as 1.5 * IQR with outliers removed.

## DISCUSSION

The last decade has witnessed an unprecedented surge in the field of RNA modifications, propelled by the advancement in mapping RNA modifications transcriptome-wide through next generation sequencing-based methods (7, 8). This has allowed researchers to establish intricate connections between RNA modifications and fundamental molecular processes such as RNA splicing, nuclear export, RNA stability and translation, unraveling previously elusive aspects of cellular regulation (63–66). Notably, several years prior to this new-found interest in epitranscriptomics, pioneering works revealed the immunosuppressive roles of certain RNA modifications, a finding that would eventually pave the way towards mRNA vaccines (67–69).

Recently, nanopore direct RNA sequencing (DRS) has solidified its position as a promising tool for investigating the epitranscriptome, due to its unique capability of sequencing native RNAs, thus making it possible to detect multiple modifications within the same native RNA molecule (17, 25, 28). Indeed, multiple works have already employed this technology to characterize the epitranscriptomic landscape of mRNAs, rRNAs and tRNAs (29, 32, 41). Early approaches for the detection of modifications using the nanopore platform have relied on the fact that modified residues will cause an error during the basecalling process (14). In this regard, several tools such as EpiNano (18), DiffErr (41), ELIGOS2 (38), DRUMMER (56) and JACUSA2 (39) have been developed to exploit this information and perform pairwise-comparisons between a modified sample and an unmodified control to identify modified sites (18, 38, 39, 41, 56). Notably, a critical component of this process is the basecaller, which is trained on a specific dataset and responsible for converting the raw current intensity signal that is output by the sequencing machine into a nucleotide sequence. The currently used basecalling model, which we refer to as ‘default’ (provided with guppy 6.0.6 and upwards), shows overall per-read accuracy of 90% using ONT’s proprietary basecaller *Guppy* (42). Previous efforts have been made to develop novel, more accurate basecalling softwares, such as RODAN for RNA and Chiron and DeepNano for DNA (70–72), among others. However, the effect of using alternative basecalling models on per-read accuracy and detection of RNA modifications has so far not been explored.

Here we present two novel basecalling models, IVT and SUP (**Table S1**), and benchmark their ability to detect a battery of RNA modification types as well as their basecalling accuracy and effect on read mapping. We find that using a model that was trained on *in vitro* transcribed RNA (IVT model) significantly improves the detection of m^6^A modifications (**Fig 1E-F**) by generating a strong deletion signature at the modified site (**Fig S2C-D**) and overall increased SumErr and ΔSumErr, leading to a significant increase in the total number of high-confidence m^6^A-sites recovered in *in vivo* samples, compared to the default model (**Fig 4B**). We argue that this improvement is due to the fact that the default model was trained on native RNA from different organisms containing large amounts of m^6^A, and it learnt to recognize both unmodified A and m^6^A as A, especially at naturally modified 5-mers (**Fig 3A**). We find evidence for this hypothesis in the fact that the most improved 5-mer coincides with the most-highly modified 5-mer in humans GGm^6^ACT (**Fig 3B, Fig S4B**).

Our second model SUP was trained with the intention of maximizing read accuracy. Therefore, we changed the model architecture from flip-flop used in both default and IVT to CTC-CRF (currently used in ONT DNA-seq, see *Methods*). We found that SUP reaches a per-read accuracy of 97% on human data compared to 91% in default, which is a substantial improvement and to our knowledge the most accurately basecalled direct RNA-sequencing run to date (**Fig 4A**). We additionally found that this improved accuracy was accompanied by an increased number of mapped reads, becoming particularly apparent in short reads, where the increase in read mappability reached ∼50% (**Fig 4B-C**). Similar performance has been reported regarding yet-to-be-released RNA-004 chemistry (96% accuracy). We should note that our improvements in basecalling accuracy (up to 97%) were obtained using the RNA-002 chemistry, suggesting that improvements in the accuracy reported for the RNA-004 chemistry are likely not related to changes in the pore chemistry but rather to improvements in the basecalling models.

## DATA AVAILABILITY

Fast5 DRS datasets basecalled with default corresponding to ‘curlcakes’ containing ac^4^C, m^5^U and m^1^Y as well as UNM-S rep2 and UNM-S rep3 (**Table S2**) have been deposited in ENA, under accession code PRJEB67632. The corresponding FASTQ files, basecalled using all three models used in this work, have been deposited in GEO, under accession code GSE246151. basecalled Fast5 datasets from unmodified ‘curlcakes’, as well as those containing m^6^A, Ψ, m^5^C and hm^5^C were taken from previously published studies (18, 25), and were already publicly available in ENA, under accession codes PRJNA511582, PRJNA549001, PRJNA563591 and PRJNA548268.

The following publicly available datasets were also used in this study: (i) raw fast5 from wildtype *and Mettl3^-/-^* HEK293T cells, used to test whether novel basecalling models can improve the detection of m^6^A *in vivo*, were downloaded from ENA (PRJEB40872) (21); (ii) raw fast5 from *M. musculus, X. laevis, S. cervisiae* and *A. thaliana* (18, 30, 41), used to compare basecalling accuracy across several model species were downloaded from SRA (SRP363295, PRJNA521324) and ENA (PRJEB32782); (iii) raw fast5 of an unmodified (UTP) and modified (m^1^ΨTP) synthetic eGFP vaccine (45), were downloaded from SRA (PRJNA856796). All project and sample accessions of all datasets used in this work are additionally listed in **Table S2**.

## CODE AVAILABILITY

All code used to build figures shown in this work can be found on GitHub (https://github.com/novoalab/basecalling_models). All trained basecalling models used in this work are available in OSF (https://osf.io/2xgkp/). Processing of raw fast5 files was performed using the MasterOfPores (version 2.0) nextflow pipeline (https://github.com/biocorecrg/MOP2) (47, 73).

## ACKNOWLEDGEMENTS

GD is part of the ROPES ITN which received funding from the European Union’s Horizon 2020 research and innovation programme under the Marie Sklodowska-Curie grant agreement no. 956810. OB is supported by funds from Merck Innovation 2020. LPP was supported by funding from the European Union’s H2020 research and innovation programme under Marie Sklodowska-Curie grant agreement No. 754422, and is currently supported by ERC funds (ERC-StG-2021 No 101042103 to EMN). JYR is supported by funds from the Swiss National Science Foundation (310030_197906) and the Deutsche Forschungsgemeinschaft (RO 4681/4-2, RO 4681/6-1, and RO 4681/12-1, TRR319 RMaP). This work was supported by funds from the Spanish Ministry of Economy, Industry and Competitiveness (MEIC) (PID2021-128193NB-100 to EMN) and the European Research Council (ERC-StG-2021 No 101042103 to EMN). We acknowledge the support of the MEIC to the EMBL partnership, Centro de Excelencia Severo Ochoa and CERCA Programme / Generalitat de Catalunya.

## AUTHOR CONTRIBUTIONS

GD performed most of the bioinformatic analyses included in this work, together with OB, SC and AD-T. LPP trained the RNA basecalling models used in this work. LLL, MCL and OB generated curlcake datasets used in this work. AC and AD-T contributed with bioinformatic analyses. OB and EMN supervised the project, with the contribution of JYR. EMN conceived the project and acquired funding. GD built the figures. GD and EMN wrote the manuscript, with the contribution from all authors.

## COMPETING INTERESTS

EMN has received travel and accommodation expenses to speak at Oxford Nanopore Technologies conferences. GD and OB have received travel bursaries from ONT to present their work at conferences. EMN is a member of the Scientific Advisory Board of IMMAGINA Biotech. The authors declare that the research was conducted in the absence of any commercial or financial relationships that could otherwise be construed as a conflict of interest.

