## Supplementary Files for "Enhanced detection of RNA modifications and mappability with high-accuracy nanopore RNA basecalling models"

* These authors contributed equally

**SUPPLEMENTARY FIGURES**

**Figure S1. Overview of benchmarked basecalling models. (A)** Schematic overview of the three basecalling models benchmarked in this study (default IVT and SUP), their training data, and the underlying architecture of their neural network **(B)** Train-loss plot for the SUP model. The training was stopped after Epoch 5. **(C)** Train-loss plot for the super accurate model (SUP) basecalling model. **(C)** Percentage of basecalled reads mapped to unmodified and fully modified 'curlcake' sequences across all tested modifications.

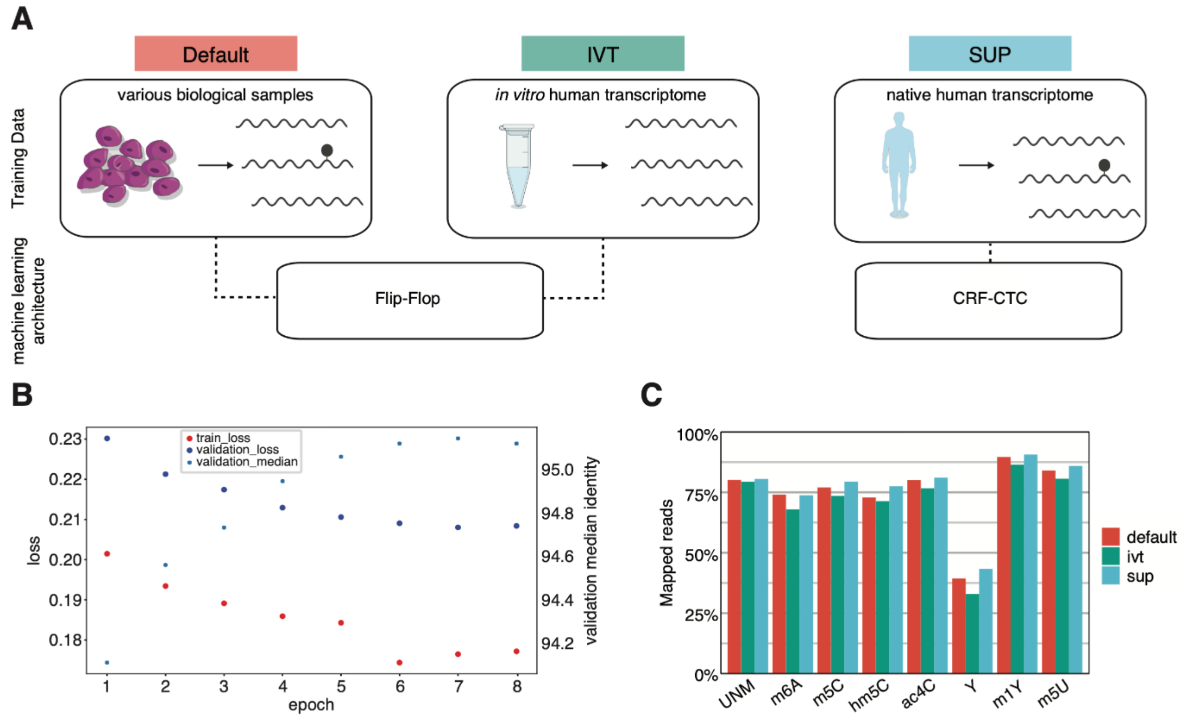

**Figure S2**. **Error Signatures of three base-calling models across a set of RNA modifications (A)** Boxplots representing the background error of each model. The summed error (mismatch-freq, deletion-freq, and insertion-freq) for every 5-mer found in the unmodified 'curlcake sequences'. Statistical analysis was performed using a two-sided Wilcoxon test with 'default' as the reference group. **(B)** Boxplots showing the delta summed error of centrally modified 5mers at position 0 for selected modifications (m$^5$U, m$^5$C, hm$^5$C, and ac$^4$C). Statistical analysis was performed using a two-sided Wilcoxon test with 'default' as the reference group. To correct for multiple-hypothesis testing the Benjamini-Hochberg procedure was used to obtain adjusted p-values. **(C)** Comparison of the median delta mismatch error (left), deletion error (middle), and insertion error (right)(deltaError = $Err_{MOD}$ - $Err_{UNM}$) for all 5-mers with a central (= position 0), modified base across the three tested base-calling models (= default, IVT and SUP). **(D)** Comparison of the median delta mismatch error (left), deletion error (middle), and insertion error (right)(= $Err_{MOD}$ - $Err_{UNM}$) for all 5-mers with a central (= position 0), modified base scaled by each modification to highlight basecaller dependent differences. For Figure S2A-B the sample size reported by *n* represents the number of 5mers contributing to each boxplot per panel. The box is limited by the lower quartile Q1 (bottom) and upper quartile Q3 (top). Whiskers are defined as 1.5 * IQR with outliers represented as individual dots. Both figures used the same p-value cutoffs (ns: p > 0.05, *: p <= 0.05, **: p <= 0.01, ***: p <= 0.001, ****: p <= 0.0001).

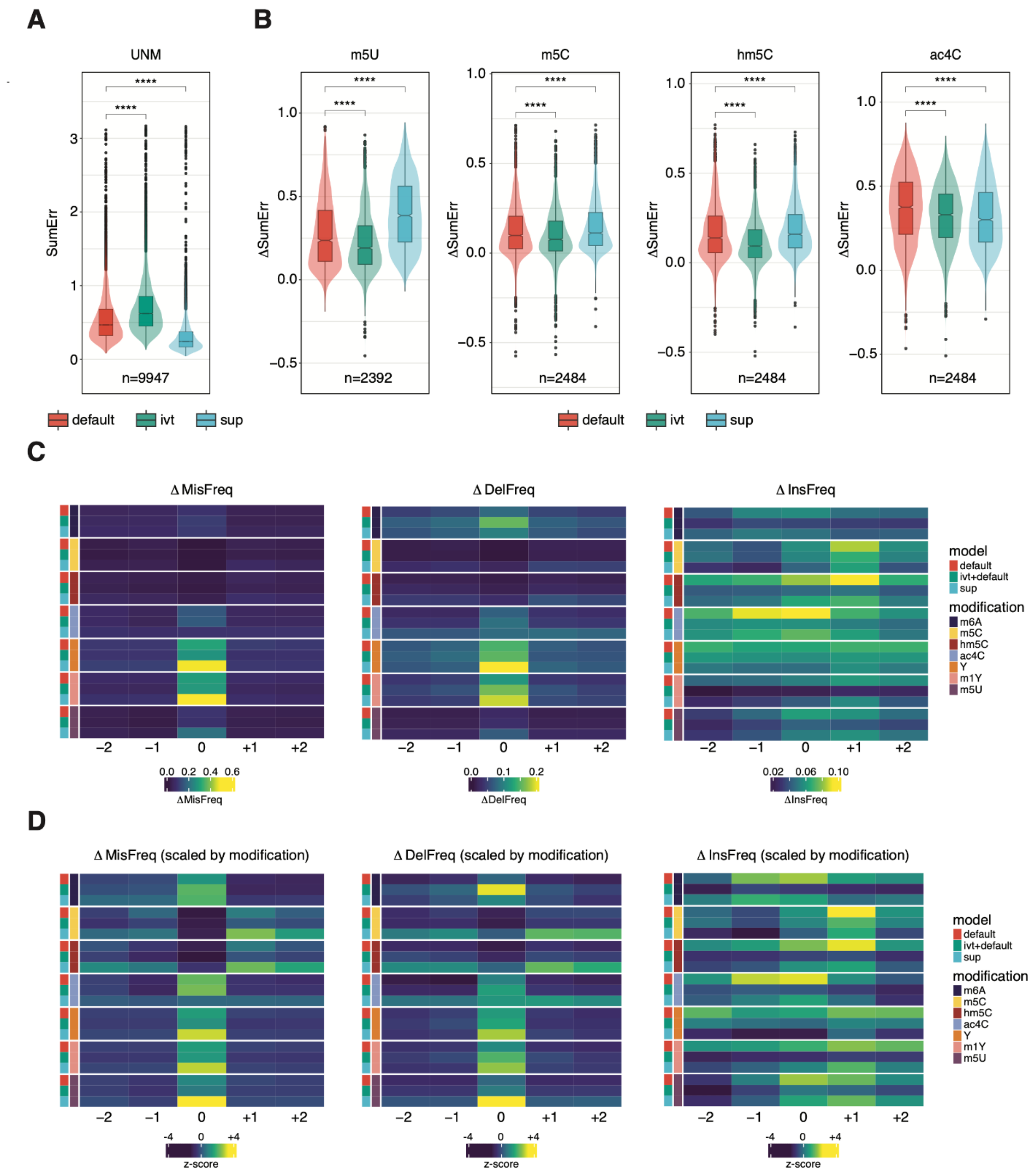

**Figure S3**. **Global mismatch error signatures across different modifications and base-calling models (A)** Global mismatch frequencies obtained from EpinanoRMS (version 1.1) per modification (rows) and base-calling model (columns) for ‘curlcakes’ 1-4 are reported. Matches to the respective reference base are colored in grey while mismatches caused by adenosine, thymine, cytidine, or guanine are colored in green, red, blue, and yellow, respectively. The percentage of matches to the reference is reported on each bar.

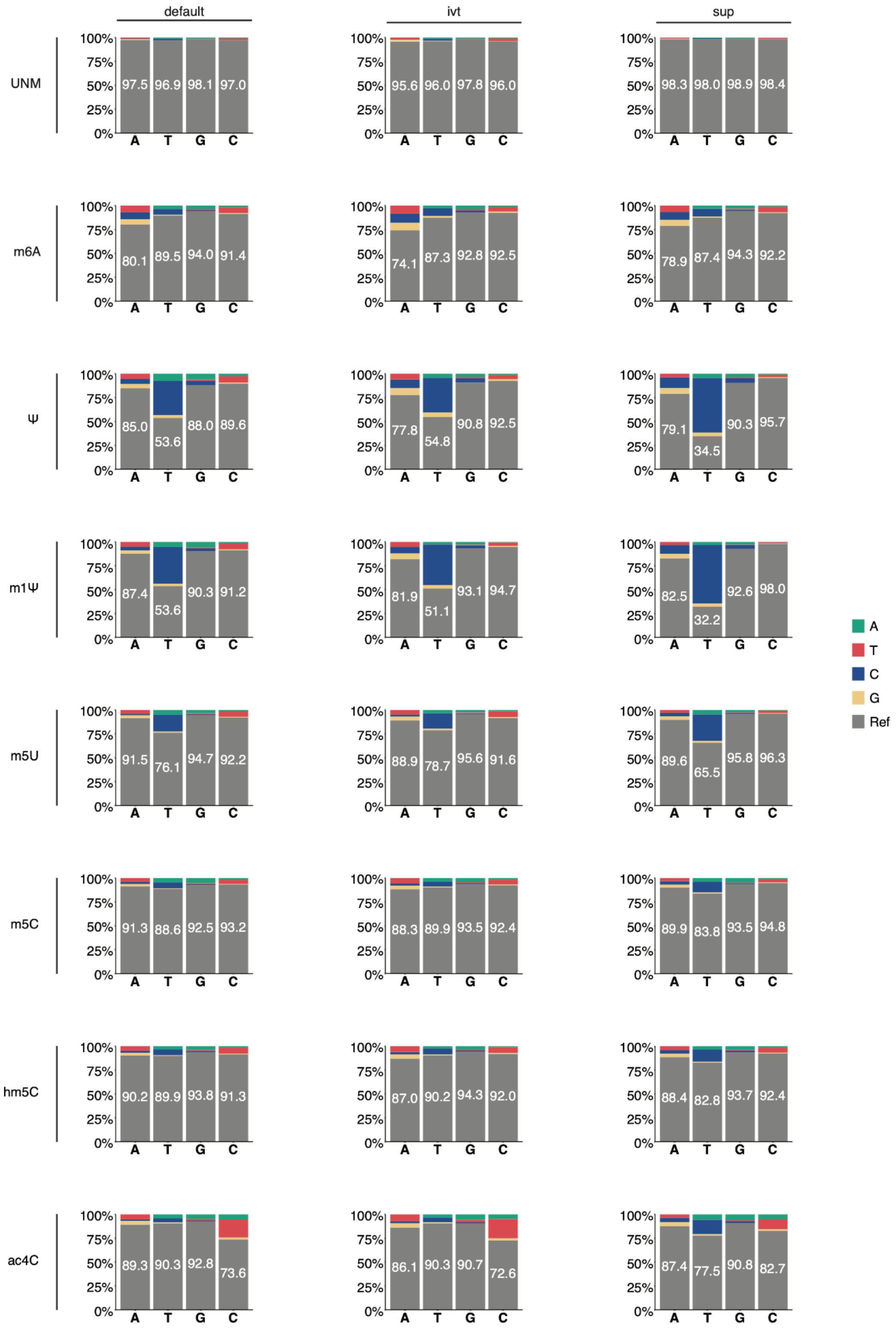

**Figure S4**. **Motif-based analysis of novel base-calling models on m⁶A modified *in vitro* transcripts. (A)** Comparison of the delta summed errors (= $SumErr_{MOD}$ - $SumErr_{UNM}$) for individual motifs considered DRACH (top) and non-DRACH (bottom) between default and SUP. Each dot represents a single observation of that 5-mer within the 'curlcake' sequences. Statistical analysis was performed using a paired two-sided Wilcoxon test. To correct for multiple-hypothesis testing the Benjamini-Hochberg procedure was used to obtain adjusted p-values (ns: p > 0.05, *: p <= 0.05, **: p <= 0.01, ***: p <= 0.001, ****: p <= 0.0001). The sample size reported by *n* represents the number of 5-mers contributing to each boxplot per panel. The box is limited by the lower quartile Q1 (bottom) and upper quartile Q3 (top). Whiskers are defined as 1.5 * IQR. **(B)** Ranking of 5-mers that contain a single, central A-position based on the amplitude of their ΔΔ summed error (= $\Delta SumErr_{IVT}$ - $\Delta SumErr_{DEFAULT}$). Whiskers are defined as 1.5 * IQR with outliers represented as individual dots. The sample size reported by *n* represents the number of individual 5-mers of that motif present in the 'curlcake' sequences.

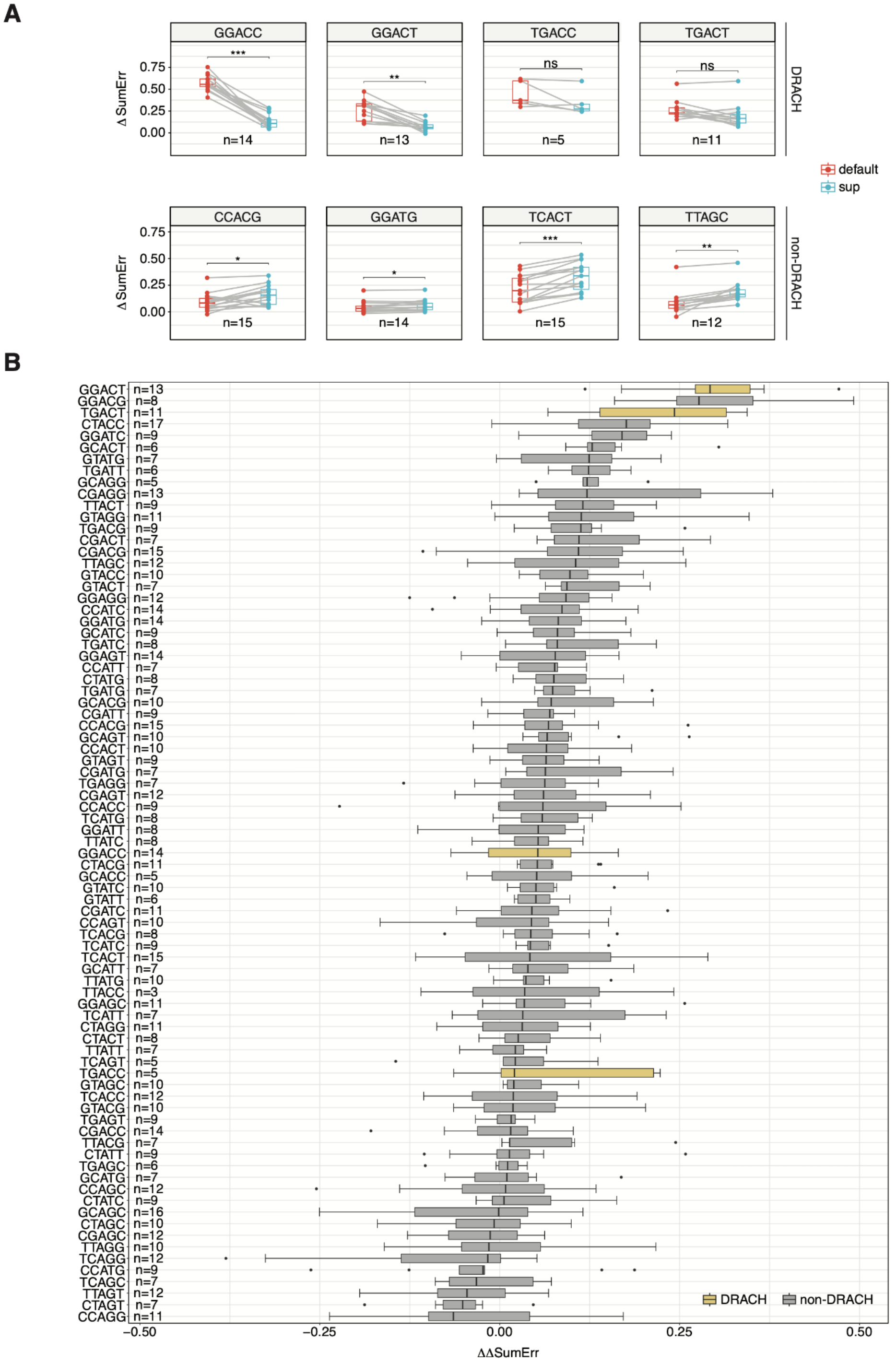

**Figure S5**. **Effect of different m$^6$A stoichiometries on the strength of error signatures across base-calling models (A)** Comparison of the error-signature amplitude (deltaSumError = $SumErr_{MOD}$ - $SumErr_{UNM}$) at different modification stoichiometries between 5mers that fall within the DRACH motif (D=A, G or U; R=G or A; H=A, C or U) and others (non-DRACH). To determine at which stoichiometry the tested models outperform the default, a one-sided non-parametric Wilcoxon test was performed with the default model as the reference group. The sample size (*n*) depicted on the side of each row corresponds to the number of samples per boxplot. Reported significance labels correspond to the BH-corrected p-values (ns: $p > 0.05$, *: $p \leq 0.05$, **: $p \leq 0.01$, ***: $p \leq 0.001$, ****: $p \leq 0.0001$). The box is limited by the lower quartile Q1 (bottom) and upper quartile Q3 (top). Whiskers are defined as 1.5 * IQR with outliers represented as individual dots. **(B)** Line plots generated by locally estimated scatterplot smoothing (loess) across a range of recorded stoichiometries for DRACH bearing motifs. Individual points represent distinct 5mers within the 'curlcake' sequences. Lightly shaded areas correspond to the 95% confidence interval.

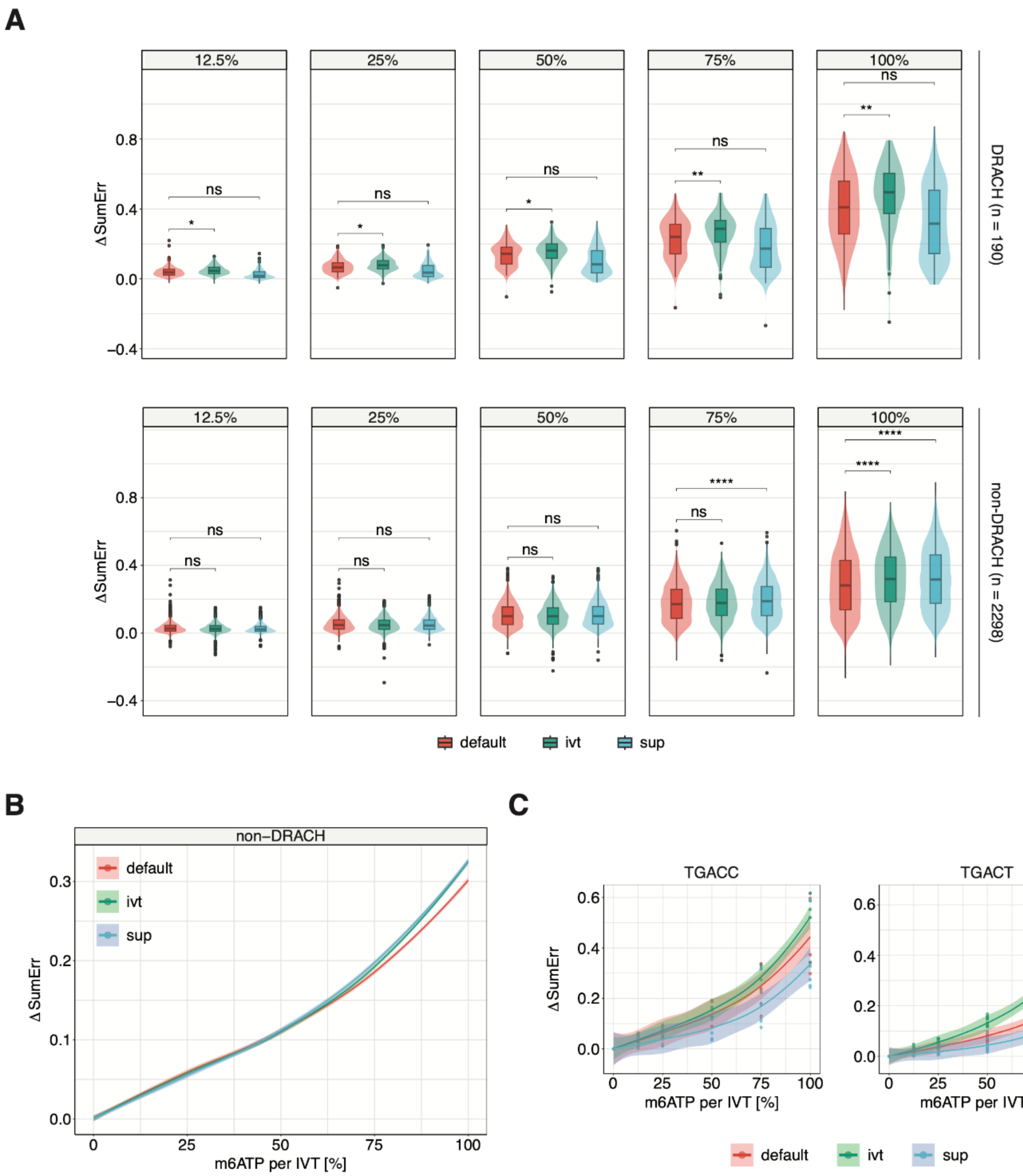

**Figure S6**. ***In-vivo* detection of m$^6$A using novel base-calling models** **(A)** Overlap of the three orthogonal m$^6$A-detection methods GLORI-seq, miCLIP, and m$^6$ACE-seq used to benchmark the *in vivo* detection ability of our basecalling models. **(B)** Overlap of coverage-matched sites between the three orthogonal methods and our SUP model **(C)** Overlap between sites detected using all three basecalling models (default, IVT and SUP). **(D)** Motif identified using BaMM for sites that did not show an overlap with at least one orthogonal method for default and IVT. **(E)** Top 5-mers reported for both models that weren't supported by at least one orthogonal method **(F)** IGV screenshot of two m$^6$A-sites *FAIM* (ENSG00000158234) and *GLRX5* (ENSG00000182512) uniquely identified by the 'IVT' model. Positions at which the mismatch frequency exceeds 0.1 are colored, while deletions are visualized as a drop in coverage.

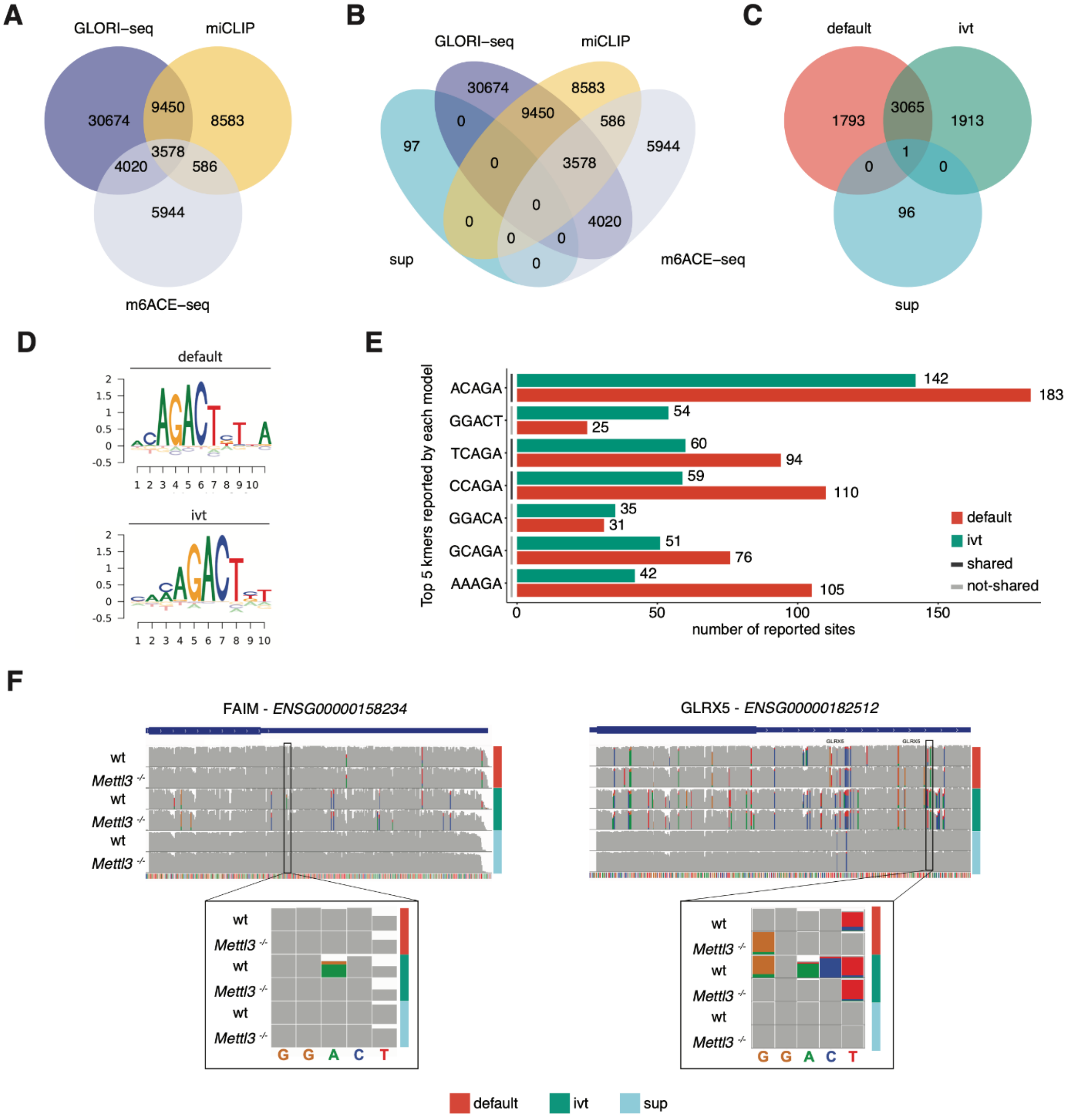

**Figure S7. Mapping comparison on different model species. (A)** Length distribution of mapped reads used to compare basecalling accuracies across the tested models. Reads were aligned to the respective reference transcriptome (Table S4). The box is limited by the lower quartile Q1 (bottom) and upper quartile Q3 (top). Whiskers are defined as 1.5 * IQR with outliers being removed. **(B)** Bars represent the amount of reads contributing to each bin used in Fig 4B to visualize differences in mapping reads caused by different basecalling models. **(C)** The relative decrease in mapped reads for IVT compared to default for both a standard mRNA library and an mRNA-short library consisting of synthetic RNA constructs with a median length of 200nt. Statistical significance was determined using a one-sided t-test, and results were corrected for multiple hypothesis testing using the Benjamini Hochberg procedure (ns: p >= 0.05, *: p < 0.05, **: p < 0.01).

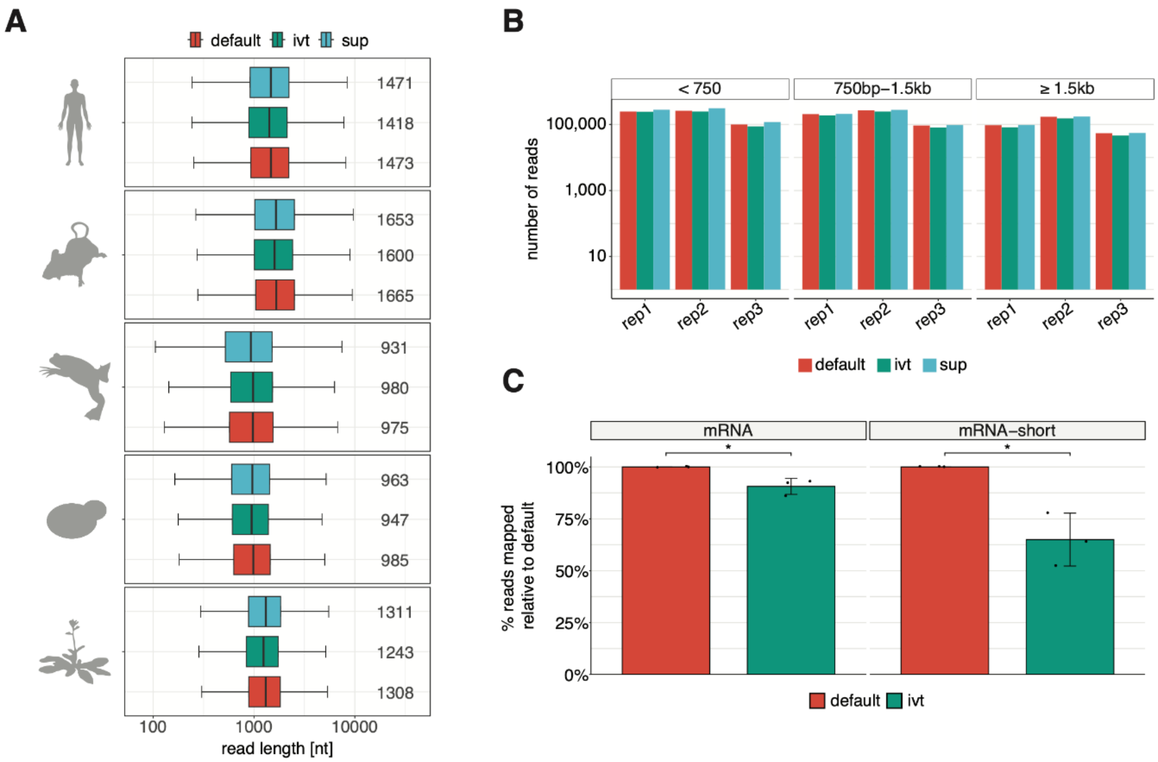

**Figure S8. Detailed inspection of error patterns caused by pseudouridine and *N1*-methylpseudouridine. (A)** Comparison of mismatch-, deletion- and insertion frequency across 5-mers that contain a single, central uridine (VVUVV). To test for statistical significance the non-parametric Wilcoxon test was used. Obtained p-values were corrected for multiple hypothesis testing using the Benjamini-Hochberg procedure. (ns: $p > 0.05$, *: $p <= 0.05$, **: $p <= 0.01$, ***: $p <= 0.001$, ****: $p <= 0.0001$). Whiskers are defined as 1.5 * IQR with outliers being removed. **(B)** Ternary plots depicting the mismatch error distribution of each model for 5-mers with a single, central uridine (VVUVV). Points are colored by the overall contribution of mismatch error to the summed error for this site (Σ mismatch-, deletion-, and insertion frequency). Barplots show the quantified mismatch error distribution for the same set of 5-mers.

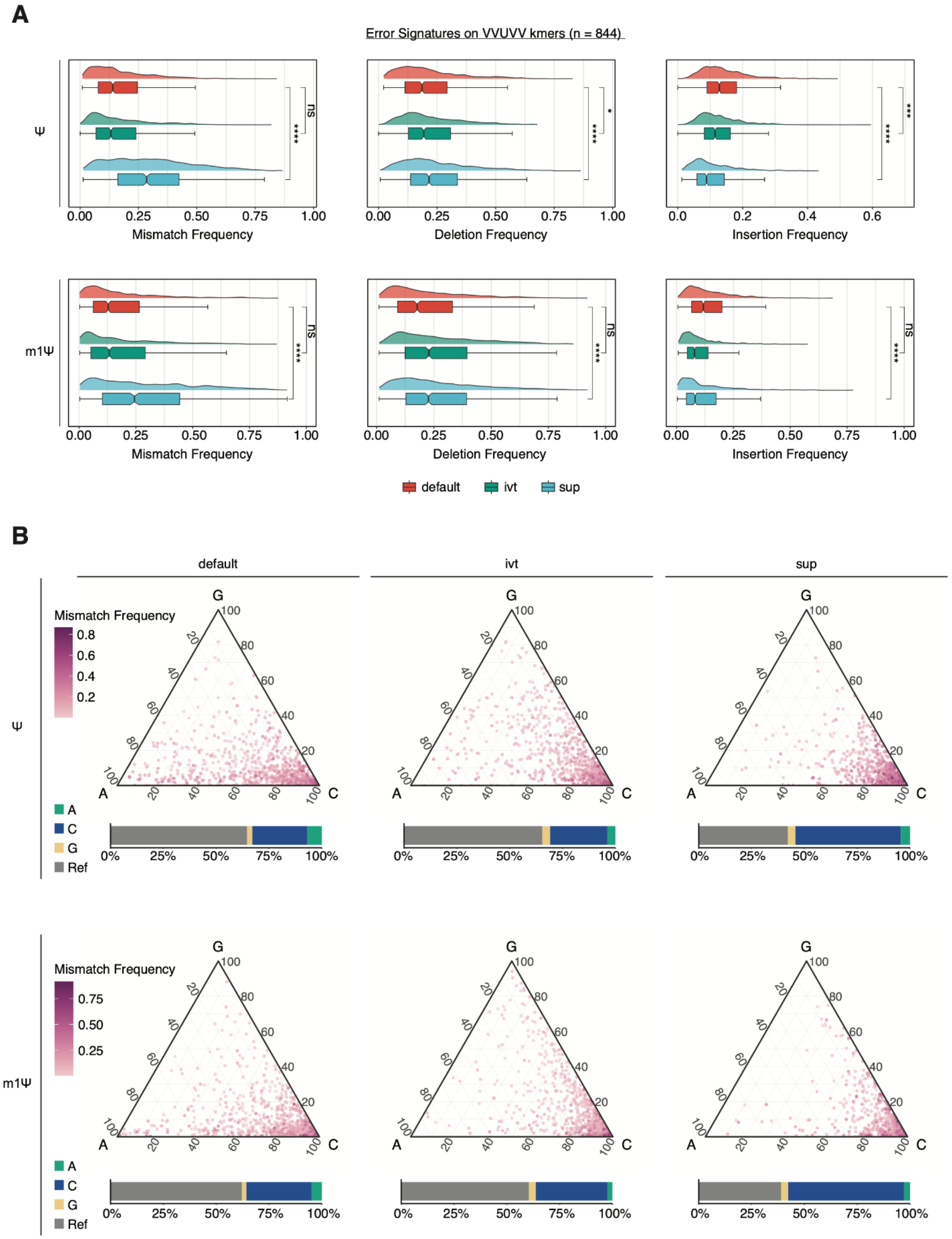

**Figure S9. Error signatures across a synthetic mRNA vaccine. (A)** Per base summed error (Σ mismatch-, deletion- and insertion frequency) for unmodified (UNM; A,C,G and U) and modifed (MOD; A,C,G an m1Ψ) eGFP mRNA vaccine. The poly-A-tail sequence was removed from this analysis to prevent artificial inflation of adenosine-based errors due to long homopolymer stretches. **(B)** Deletion- and insertion frequency at m1Ψ positions (n = 170). To test for statistical significance the non-parametric Wilcoxon test was used and values corrected for multiple hypothesis testing using the Benjamini-Hochberg procedure. (ns: p > 0.05, *: p <= 0.05, **: p <= 0.01, ***: p <= 0.001, ****: p <= 0.0001). **(C)** Comparison of global U>C conversion between default and sup. The median is indicated as a dashed line (n = 170). **(D)** Comparison of the overall summed error (Σ mismatch-, deletion-, and, insertion frequency) for sites shown in Fig 5D. For Figures 9A, 9B, and 9D the box is limited by the lower quartile Q1 (bottom) and upper quartile Q3 (top). Whiskers are defined as 1.5 * IQR with outliers removed.

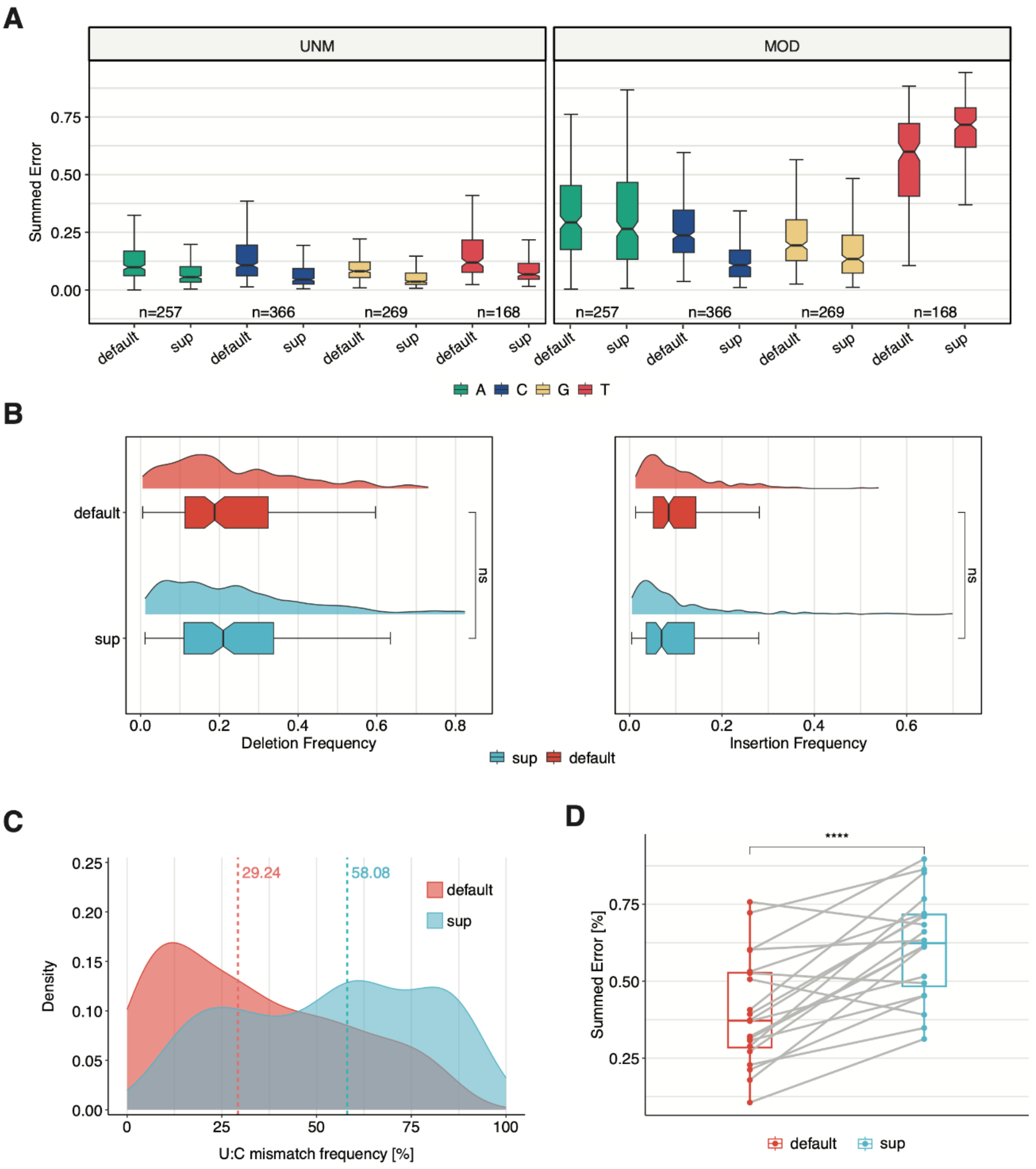
